# Increased expression of schizophrenia-associated gene C4 leads to hypoconnectivity of prefrontal cortex and reduced social interaction

**DOI:** 10.1101/598342

**Authors:** Ashley L. Comer, Tushare Jinadasa, Lisa N. Kretsge, Thanh P.H. Nguyen, Jungjoon Lee, Elena R. Newmark, Frances S. Hausmann, SaraAnn Rosenthal, Kevin Liu Kot, William W. Yen, Alberto Cruz-Martín

## Abstract

Schizophrenia is a severe mental disorder with an unclear pathophysiology. Increased expression of the immune gene C4 has been linked to a greater risk of developing schizophrenia; however, it is unknown whether C4 plays a causative role in this brain disorder. Using confocal imaging and whole-cell electrophysiology, we demonstrate that overexpression of C4 in mouse prefrontal cortex neurons leads to perturbations in dendritic spine development and hypoconnectivity, which mirror neuropathologies found in schizophrenia. We find evidence that microglia-neuron interactions and microglia-mediated synaptic engulfment are enhanced with increased expression of C4. We also show that C4-dependent circuit dysfunction in the frontal cortex leads to decreased social interactions in juvenile mice. These results demonstrate that increased expression of the schizophrenia-associated gene C4 causes aberrant circuit wiring in the developing prefrontal cortex and leads to deficits in early social behavior, suggesting that altered C4 expression contributes directly to schizophrenia pathogenesis.

## INTRODUCTION

Schizophrenia (SCZ) is a neurodevelopmental disorder characterized by disruptions in brain connectivity that lead to an array of symptoms including psychosis and deficits in cognition and social interactions (Kahn et al., 2015; Millan et al., 2016). Current treatment options are fairly effective for treating psychosis, but do not improve cognitive or social deficits, both of which contribute more significantly to the long-term prognosis of SCZ patients (Lieberman et al., 2005; Green et al., 2015; Green, 2016). Social deficits are often present in children and adolescents before the onset of psychosis, thus making early social deficits a feature of the disease (Velthorst et al., 2017). Therefore, understanding the underlying mechanisms of early social deficits in SCZ could open a therapeutic window to alter disease progression in these individuals.

Among the many clinical traits of SCZ are deficits in social cognition such as social cue perception and emotion regulation, which suggest dysfunction of the prefrontal cortex (PFC) (Green et al., 2015). Moreover, patients with SCZ exhibit abnormal activity in the PFC during affective face perception and cognitive reappraisal (Morris et al., 2012; Taylor et al., 2012; Delvecchio et al., 2013). These results are not surprising given the well-established role of the PFC in social behaviors (Yizhar, 2012, Ko, 2017, Grossmann, 2013). Although the mechanisms that cause social deficits in SCZ are unclear, evidence suggests that dysfunction in the PFC correlates with symptom onset and severity (Vita et al., 2012). In fact, a neurological hallmark of SCZ is a reduction of gray matter in the PFC due to a loss of neuronal processes and synapses (Glantz and Lewis, 2000; Thompson et al., 2001). Therefore, it has been hypothesized that pathology in SCZ arises in part due to deficits in the pruning of cortical synapses, thus producing aberrant circuitry.

In support of this, recent studies have found that a highly-associated SCZ gene involved in innate immunity, complement component 4 (C4), is involved in the developmental refinement of retinogeniculate synapses (Sekar et al., 2016). C4 is a member of a cascade called the complement system that, in the innate immune system, forms a proteolytic protein cascade that clears debris, enhances inflammation, and tags pathogens for engulfment or destruction (Nesargikar et al., 2012). The classic complement cascade is activated by an initiator protein called complement component 1 subunit q (C1q), which activates downstream proteins by proteolytic cleavage. Once the final complement protein in this cascade, complement component 3 (C3), is activated by cleavage, it binds to the pathogen and is recognized and phagocytosed by macrophages that express the complement component 3 receptor (CR3) (Sarma and Ward, 2011). It has been hypothesized that the complement cascade’s molecular machinery used to eliminate pathogens in the periphery also tags synapses for elimination in the brain (Stevens et al., 2007). It has been shown that microglia, the brain’s resident macrophage, are able to engulf neuronal and synaptic material dependent on the presence of complement proteins (Schafer et al, 2012). Microglia are the only cells in the brain that express CR3, which allows microglia to recognize and phagocytose material tagged by the complement cascade (Zabel and Kirsch, 2013). It has also been found that microglia-mediated synaptic pruning is necessary for normal development of the brain, but can contribute to pathology when mis-regulated by causing abnormal wiring of circuits (Paolicelli et al., 2011).

Despite significant progress in this field, it is still not clear how reciprocal interactions between neurons and microglia contribute to the maturation and refinement of developing cortical circuits (Tremblay et al., 2010; Hoshiko et al., 2012; Miyamoto et al., 2016). Mouse studies investigating the role of C4 and other complement proteins in synaptic refinement have relied on loss of function manipulations (Stevens et al., 2007; Schafer et al., 2012; Martinelli et al., 2016), although SCZ is linked to increased C4 expression (Sekar et al., 2016). Additionally, previous studies have focused on the role of complement proteins in the visual system rather than interrogating their role in SCZ-relevant brain regions, such as the PFC. Thus, overall, it is not known how complement protein expression impacts the development of cortical microcircuits, which are commonly altered in neurodevelopmental disorders (Meredith, 2015; Del Pino et al., 2018). A fundamental question that remains unanswered is whether increased expression of C4 causes deficits in PFC circuitry and SCZ-relevant behaviors.

To better understand the role of C4 in the pathogenesis of SCZ, we studied the impact of increased expression of C4 in cortical regions relevant to this disease (Glantz and Lewis, 2000). Here, we investigated the effects of C4 overexpression on developing networks in layer (L) 2/3 of the mouse medial prefrontal cortex (mPFC), a cortical region where gray matter loss is most prominent in SCZ (Barch et al., 2001; Beneyto and Lewis, 2011). We show that C4 overexpression induced transient structural and functional changes to L2/3 excitatory neurons in the mPFC. Additionally, we interrogated the role of microglia in complement-induced wiring deficits in the mPFC by measuring microglia-neuron interactions and microglia engulfment of synaptic material. Lastly, we examined the relationship between C4-induced PFC circuit dysfunction and abnormalities in rodent social behavior. Taken together, we link C4 expression to neural and behavioral deficits relevant to SCZ, and identify a critical window where the trajectory of brain development might be able to be therapeutically targeted in SCZ.

## RESULTS

### C4 is expressed in neurons of the medial prefrontal cortex and can be overexpressed using *in utero* electroporation

In initial experiments using *in situ* hybridization, we showed that PFC neurons in postnatal day (P) 30 control mice express low levels of *C4B* transcript (Figure 1). C4B transcript was not present in tissue from *C4B* knock-out (C4 KO) mice (Supplemental Figure 1), which demonstrates the specificity of the mouse C4B (mC4) *in situ* hybridization probe. To target layer (L) 2/3 mPFC pyramidal neurons for C4 overexpression, we used *in utero* electroporation (IUE) in mice at embryonic day (E) 16 (Szczurkowska et al., 2016) (Figure 1A and B). Human C4 (hC4) contains two distinct genes, or isotypes, C4A and C4B (Belt et al., 1985); whereas mouse C4 (mC4), or C4B, contains an amino acid sequence that is highly similar to both of the two human C4 isotypes (Nonaka et al., 1985). We performed IUE using plasmids with the CAG promoter to co-express GFP and mouse *C4B* (mC4 condition) (Figure 1B and C). Then, we used multiplex *in situ* hybridization at P21 to quantify the extent of C4 overexpression achieved via IUE delivery. Within the same coronal section of mPFC, we identified cells that were transfected (GFP+/CaMKIIα+) and untransfected (GFP-/CaMKIIα+), and compared the percent of their cell bodies that were also positive for C4 mRNA (Cell body area C4+/Total cell body area) (Figure 1C and D). This allowed us to quantify both endogenous expression of mC4 in pyramidal neurons and the extent of mC4 overexpression in transfected cells.

**Figure 1:**
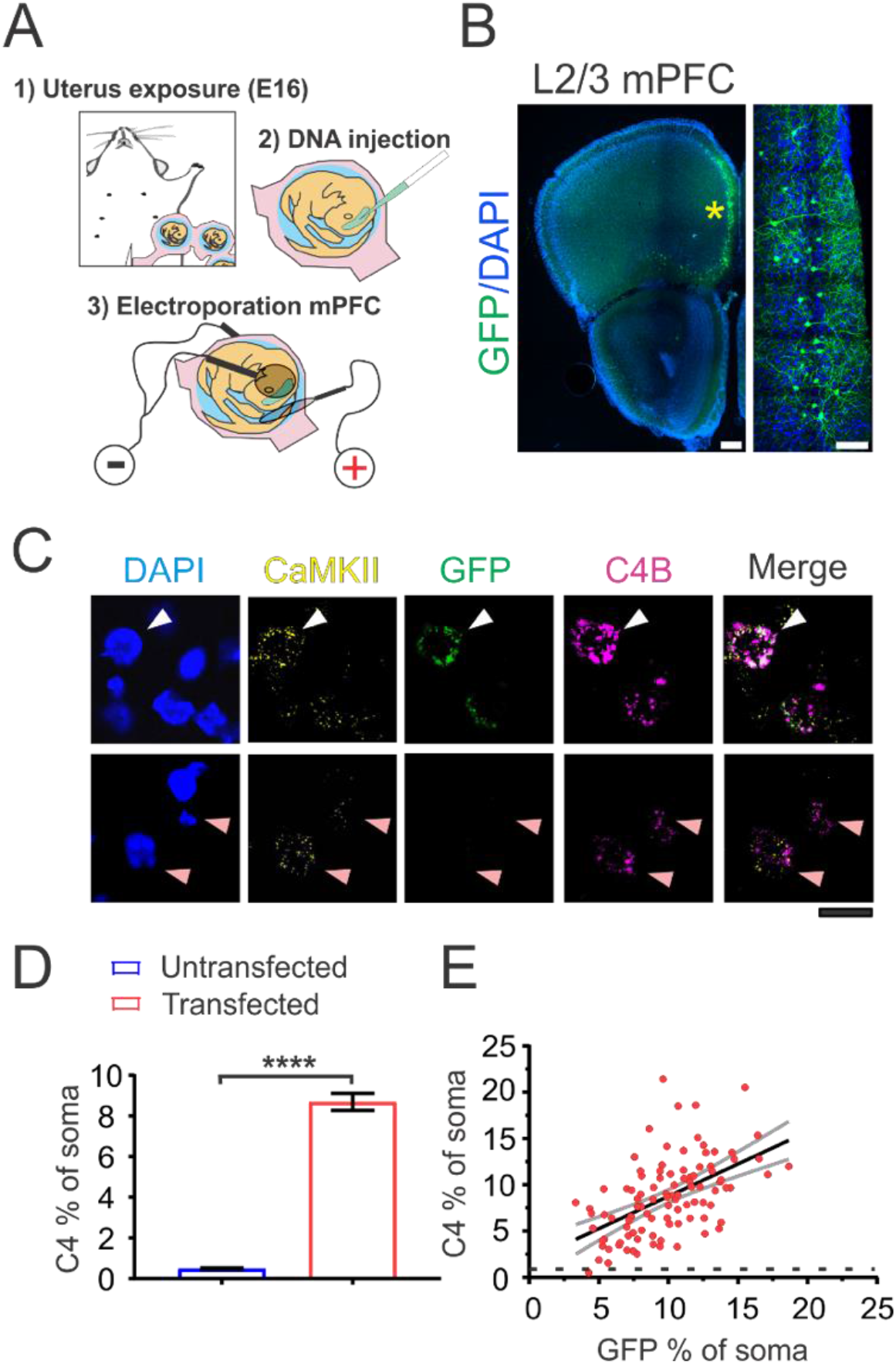
C4 is expressed in neurons of the medial prefrontal cortex and can be overexpressed using *in utero* electroporation. **(A)** Diagram depicting IUE surgery performed in E16 dams. **(B)** Representative 20X confocal image of IUE with GFP targeted to L2/3 mPFC. Yellow asterisk: L2/3 GFP+ neurons. Left panel scale bar = 250 μm. Right panel scale bar = 75 μm. **(C)** Representative 60X confocal images of *in situ* hybridization from the same coronal section showing CaMKIIα+ neurons that are transfected with GFP and mC4 (C4B) (white arrowhead) and untransfected neighbors expressing mC4 (pink arrowhead) in P21 mice. Scale bar = 15 μm. **(D)** IUE reliably increases C4B transcript level in transfected cells. Percent of soma area positive for C4 transcript in transfected and untransfected neurons. N = 100 neurons (3 mice) per condition. t-test. \*\*\*\**p* < 0.0001. Mean and SEM. **(E)** Transcript levels of GFP and mC4 positively correlate in transfected cells. Black line: linear fit. Gray lines: 95% confidence intervals. Black dotted line: average endogenous C4 expression in CaMKIIα+ mPFC L2/3 neurons. N = 100 transfected neurons (3 mice). Pearson’s R correlation and linear regression. R^2^ = 0.28. \*\*\*\**p* < 0.0001.

IUEs reliably increased expression of mC4 in excitatory CaMKIIα+ neurons of the mPFC by 17.8-fold relative to their nearby untransfected neighbors (Figure 1C and D, t-test, \*\*\*\**p* < 0.0001). By comparing mC4 transcript levels in transfected neurons to baseline expression in untransfected neighbors in the same tissue, we controlled for differences in mC4 transcript between individual mice. There was also a positive correlation between amount of GFP and mC4 transcript present in transfected cells (Figure 1E), indicating that neurons with similar GFP fluorescence also had similar levels of mC4 mRNA. Furthermore, we found that 99% of cells expressing GFP had higher amounts of mC4 mRNA relative to the mean transcript levels of nearby untransfected cells (Figure 1D and E). Overall, these results show that mPFC L2/3 neurons express C4 and that we can overexpress C4 in these cells using IUE.

### mC4 overexpression causes dendritic spine alterations in apical arbors of L2/3 neurons

Dendritic spines are small membranous protrusions that emerge from the dendrites of neurons and mediate the vast majority of excitatory synaptic transmission in the cortex (Rochefort and Konnerth, 2012). The morphology and density of these postsynaptic structures change throughout development, reflecting modifications in the maturation and strength of excitatory connections (Portera-Cailliau et al., 2003; Forrest et al., 2018). To test whether increased expression of C4 in developing neurons leads to structural alterations in dendritic spines, we used IUE to produce control conditions (pCAG-GFP) or C4 overexpression conditions (pCAG-GFP with pCAG-mC4B or pCAG-hC4A, for mC4 and hC4 conditions, respectively). Since early postnatal development is an important period for synaptic maturation and refinement of cortical circuitry (Cohen-Cory, 2002), we measure dendritic spine density using confocal imaging of GFP in fixed brain sections collected at P7-9, P14-16, P21-23 and P55-60.

In control conditions, the density of apical tuft dendriti c spines was developmentally regulated and increased nearly eightfold in the two weeks from P7 to P21 (Figure 2A, Tukey’s test, \*\*\*\**p* < 0.0001). However, the developmental increase in apical tuft dendritic spines was ~30% lower in P21 neurons overexpressing mC4 or hC4 (Figure 2A and B). There were no differences in apical tuft protrusion density between any groups at P60 (Figure 2A). These results suggest that overexpression of C4 alters the developmental maturation of dendritic spines and that the human and mouse homologs of C4 are both able to cause spine dysgenesis. Overexpression of the mouse homologue of C4 led to the most dramatic defects in dendritic spine density, therefore in subsequent experiments we manipulated levels of mC4 to investigate its role in cortical circuit function.

**Figure 2:**
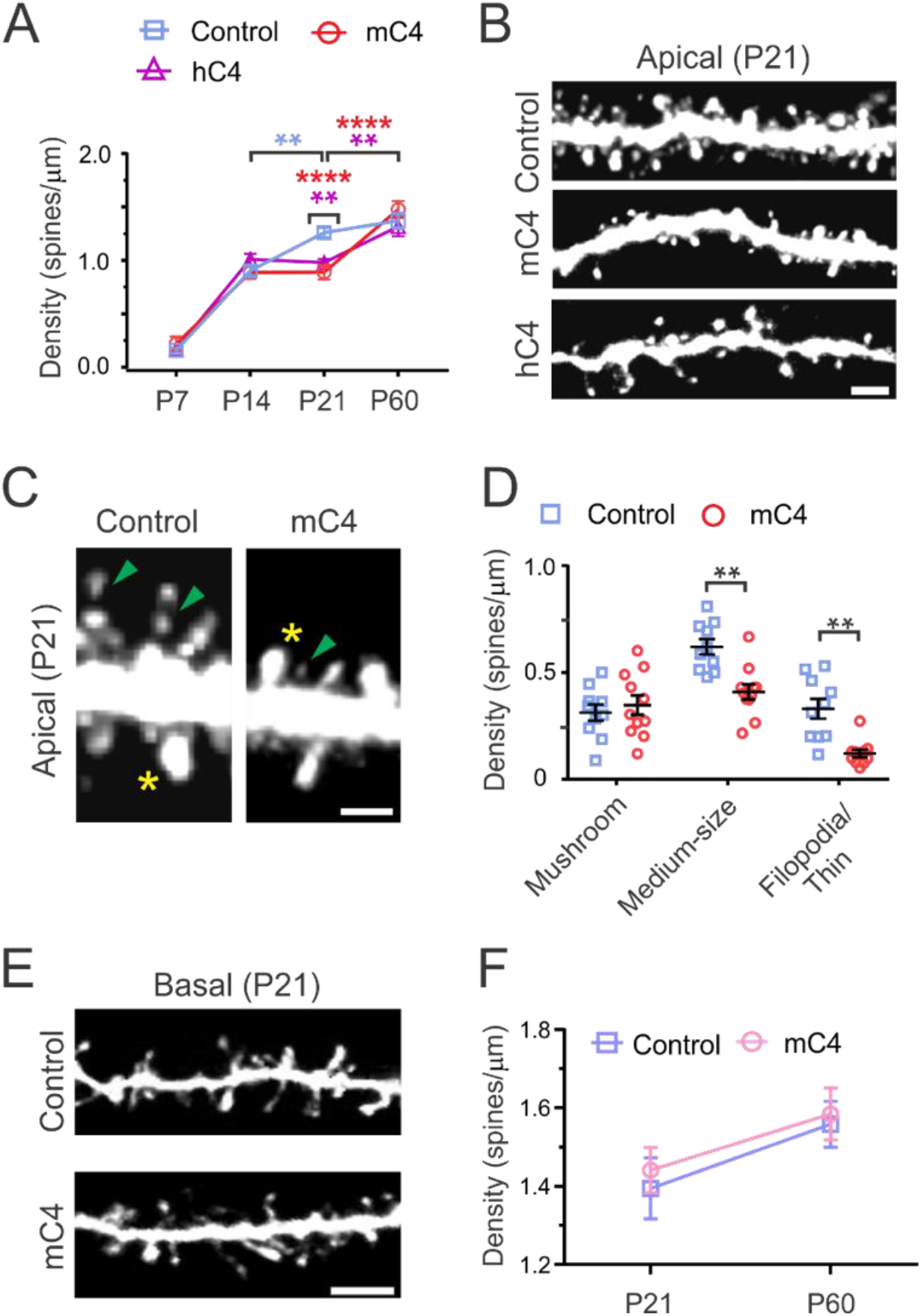
mC4 overexpression leads to dendritic spine alterations in apical arbors of L2/3 mPFC neurons. **(A)** Developmental time course reveals significant decrease of spine density in neurons overexpressing C4 at P21-23. \*\**p* < 0.01, \*\*\*\**p* < 0.0001. (B) Representative 40X confocal images of P21-23 apical dendritic tufts. Scale bar = 5 μm. **(C)** Representative 40X confocal images of P21-23 apical dendritic spine types. Yellow asterisk: large mushroom spine (TIB (a.u.) > 75%). Green arrowhead: thin spine/filopodia (TIB (a.u.) < 25%). Scale bar = 3 μm. **(D)** Spine density sorted by spine types (spine/μm) reveals a specific <u>reduction</u> of medium-sized and thin/filopodia spine types in the mC4 condition. \*\**p* < 0.01. **(E)** Representative 40X confocal images of P21-23 basal dendritic spines. Scale bar = 5 μm. **(F)** Analysis of basilar spine density (spine/μm) reveals no difference across groups. **(A-F)** N = 10-12 dendrites (from 4 mice) for each condition. Two-way ANOVA and Tukey’s test. Control: blue, mC4: red, hC4: purple. Mean and SEM.

Closer inspection of protrusion morphological types at P21 revealed a significant reduction of thin spines/filopodia and medium-sized spines in mC4-overexpressing neurons compared to controls, while putative mushroom spines remained intact (Figure 2C and D; spine types sorted by total integrated brightness, see methods). Since dendritic spine volume positively correlates with postsynaptic density size and synaptic strength (Arellano et al., 2007; Holtmaat and Svoboda, 2009), this result suggests that increased expression of mC4 preferentially altered the development of weaker synaptic connections.

We also performed morphological analysis of dendritic spines in the basal dendrites of L2/3 mPFC neurons. In contrast to the structural alterations we observed in apical tufts, increased expression of mC4 did not significantly alter the density of spines in basal dendrites compared to control conditions (Figure 2E and F). Taken together, these results demonstrate that increased expression of C4 in developing cortical neurons causes input-specific dendritic spine pathology and that less mature protrusions are targeted while larger mushroom spines remain intact.

### C4 overexpression reduces functional connectivity in cortical neurons

To determine whether the morphological spine deficits observed in C4-overexpressing cells were accompanied by functional connectivity changes, we performed electrophysiological whole-cell voltage clamp recordings in acute brain slices prepared from mPFC of P18-25 control and mC4 conditions. Voltage-clamp recordings demonstrated that overexpression of mC4 caused a decrease in the frequency of miniature excitatory postsynaptic currents (mEPSCs) evident through a ~60% decrease in mEPSC frequency and an increase in the interevent interval (IEI) (Figure 3A-C). Together with the reduction in spine density observed at P21 (Figure 2A), these results suggest that mC4 regulates excitatory synapse connectivity of L2/3 mPFC neurons by decreasing synapse density, which would lead to a decrease in mEPSC frequency. Increased levels of mC4 also caused a ~18% reduction in the amplitude of mEPSCs (Figure 3D and E), demonstrating that in addition to controlling the density of synaptic connections, mC4 also modifies the postsynaptic responses of L2/3 neurons. Further analysis of postsynaptic current kinetics revealed no differences in mEPSC Rise_10-90_ or Decay_tau_ between control and mC4-transfected neurons (Figure 3F and G). Although overexpression of mC4 did not alter the frequency or the amplitude of miniature inhibitory postsynaptic currents (Figure 3H-L), it significantly slowed their kinetics (Figure 3M and N, 27% and 19% increase in Rise_10-90_ and Decay_tau_, respectively), suggesting C4 dependent changes in inhibitory synaptic location relative to the soma. Taken together, our data suggest that increased expression of C4 alters the developmental wiring of cortical neurons by decreasing their excitatory synaptic drive.

**Figure 3:**
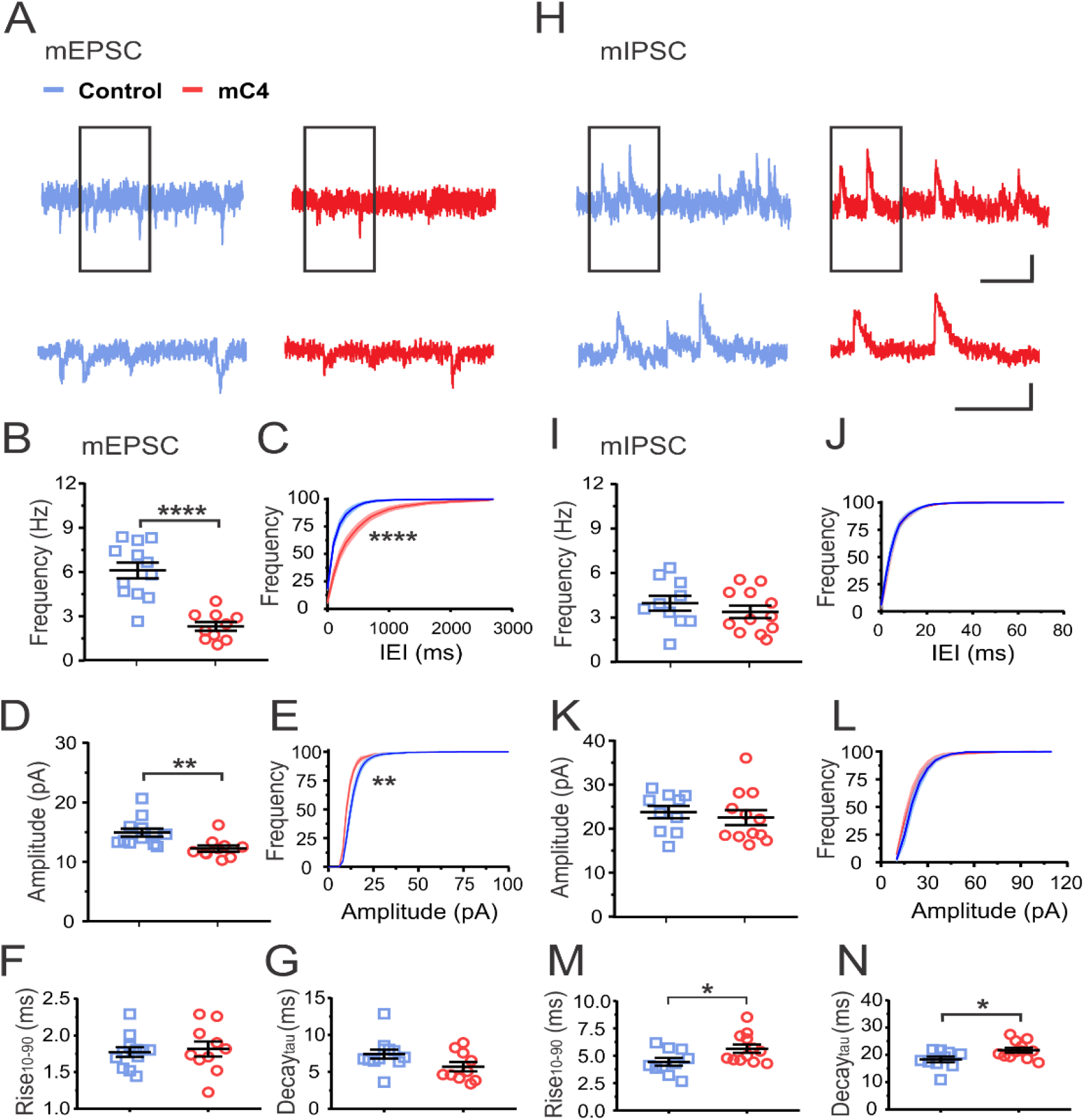
C4 overexpression reduces functional connectivity in cortical neurons. **(A)** Top: Representative whole-cell voltage clamp recordings showing mEPSCs. Top scale bar = 250 ms/10 pA. Bottom: same as top traces but expanded (black rectangle region). Bottom scale bar = 125 ms/10 pA. **(B)** Increased mC4 expression causes a reduction in mEPSC frequency. t-test. \*\*\*\**p* < 0.0001. **(C)** Overexpression of mC4 causes a rightward shift in the distribution of mEPSC interevent interval (IEI). Kolmogorov-Smirnov test. \*\*\*\**p* < 0.0001. **(D)** mC4 causes a reduction in mEPSC amplitude. t-test. \*\**p* < 0.01. **(E)** mC4 overexpression causes a leftward shift in the mEPSC amplitude distribution. Kolmogorov-Smirnov test. \*\**p* < 0.01. **(F)** mEPSC Rise_10-90_ is not altered by mC4 overexpression. t-test. *p* = 0.715. **(G)** No changes in mEPSC Decay_tau_ with mC4 overexpression. t-test. *p* = 0.07. **(B-G)** N = 12 control neurons, N = 10 mC4 neurons. Mean and SEM. Control: blue, mC4: red. **(H)** Top: Representative recordings showing mIPSCs. Top scale bar = 250 ms/20 pA. Bottom traces are same as **(H)** top but expanded (rectangle region). Top scale bar = 125 ms/20 pA. **(I)** No difference in mIPSC frequency between control and mC4 conditions. t-test. *p* = 0.3726. **(J)** Distribution of mIPSC IEIs is not changed by increased expression of mC4. Kolmogorov-Smirnov test. *p* > 0.05. **(K)** No changes in mIPSC amplitude with increased expression of mC4. t-test. *p* = 0.5832. (L) mIPSC amplitude distribution is not changed by increased expression of mC4. Kolmogorov-Smirnov test. *p* > 0.05. **(M)** mIPSC Rise_10-90_ is increased in neurons overexpressing mC4. t-test. \**p* = 0.0329. **(N)** mIPSC Decay_tau_ is increased in neurons overexpressing mC4. t-test. \**p* = 0.0225. **(I-N)** N = 10 control neurons, N = 12 mC4 neurons. Control: blue, mC4: red. Mean and SEM.

### Overexpression of mC4 decreases membrane capacitance without altering overall excitability

We also monitored the active electrophysiological properties of L2/3 mPFC neurons to determine whether overexpression of mC4 altered neuronal excitability. To do this, we injected hyperpolarizing and depolarizing current step injections of different amplitudes and recorded changes in membrane potential (V_m_) in control and mC4-OX neurons (Figure 4A and B). Overexpression of mC4 did not alter the number of action potentials (APs) in response to step current injections of incrementally increasing amplitudes (Figure 4C). In agreement with this, we did not see a change in the interevent interval of APs in response to step current injections (Figure 4D) or the minimal current injection that results in AP generation (Figure 4E) in neurons overexpressing mC4. We did not observe any complement-dependent changes in AP amplitude, AP Rise_10-90_, or AP Decay_10-90_ (data not shown). Together, these results suggest that overexpression of mC4 does not alter the intrinsic excitability of cortical neurons. In contrast, increased levels of mC4 lead to ~29% decrease in the membranecapacitance (C_m_) relative to controls (Figure 4F). This effect in passive membrane properties was specific because increased expression of mC4 did not alter the neuron’s resting membrane potential or membrane input resistance (Figure 4G and H). Since the C_m_ is proportional to the cell’s surface area, this result suggests that mC4 affects the gross morphology of cortical neurons or plasma membrane properties. Further investigation of neuronal morphology showed no difference between conditions in cell body area/diameter or in the width of primary dendrites (Supplemental Figure 2); there was also no change in dendritic complexity (Supplemental Figure 3), suggesting that gross neuronal morphology is not affected by mC4 overexpression. Besides the complement-dependent changes in C_m_, increased expression of mC4 did not have an effect on the morphology or intrinsic properties of transfected cells, thus suggesting that overall neuronal health is not affected by increased expression of mC4.

**Figure 4:**
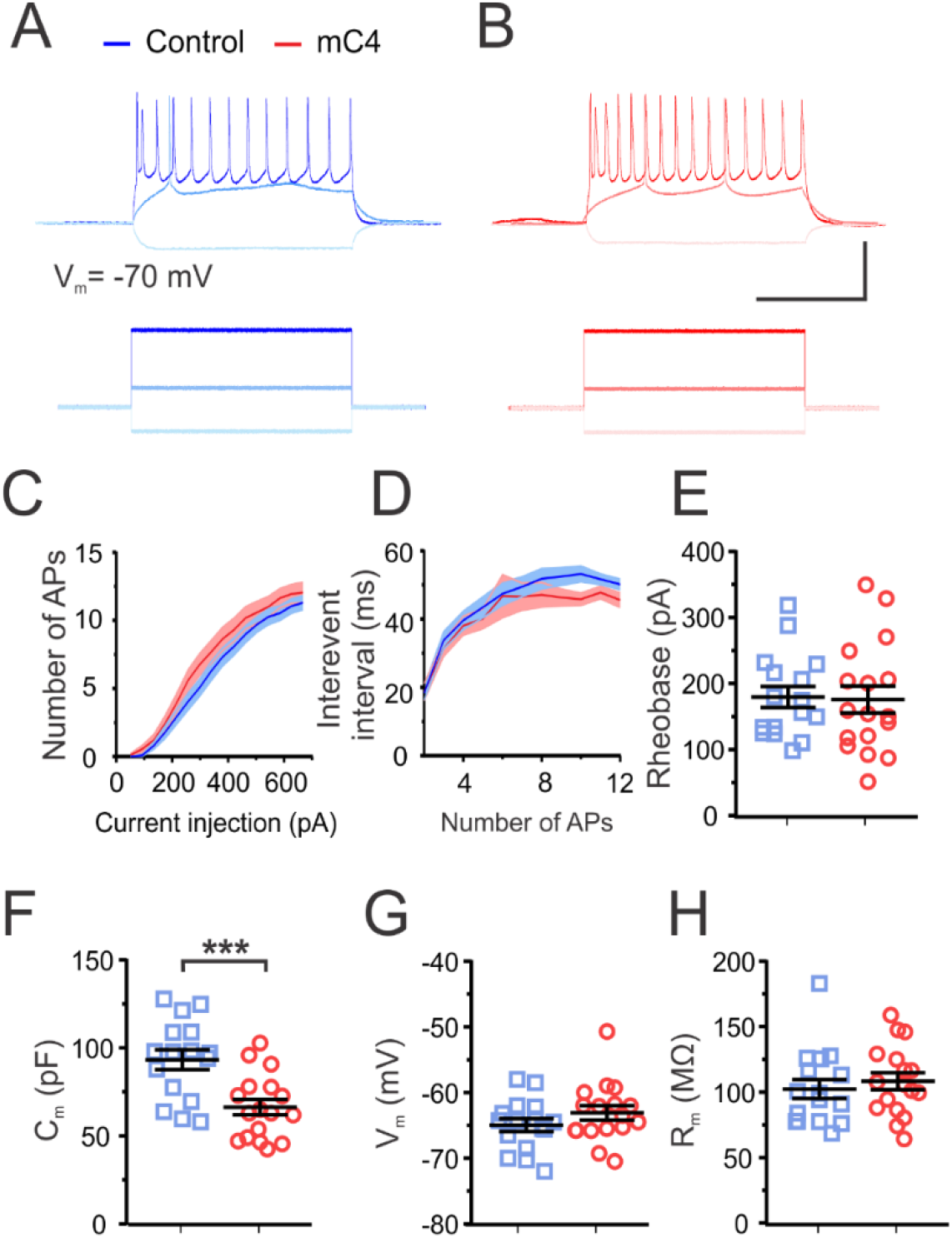
Overexpression of mC4 decreases membrane capacitance without altering overall excitability. **(A-B)** Representative current clamp recordings from control (A) and mC4 (B) neurons in response to injection of constant-current pulses. Voltage traces (top) shown for response to −220 pA (light blue/pink), 185 pA (medium blue/red) and 685 pA (dark blue/red) current injections (bottom). Scale bar = 500 pA or 50 mV. Scale bar = 250 ms. **(C)** The number of APs were not different between conditions. Dark blue trace: control mean. Light blue trace: control SEM. Dark red trace: mC4 mean. Light red trace: mC4 SEM. Two-way ANOVA. *p* = 0.9989. **(D)** Interevent interval was not different between conditions. Dark blue trace: control mean. Light blue trace: control SEM. Dark red trace: mC4 mean. Light red trace: mC4 SEM. Two-way ANOVA. *p* = 0.9838. **(E)** Rheobase was not altered by the overexpression of mC4. t-test. *p* = 0.8795. **(F)** mC4 overexpression led to a dramatic reduction in C_m_. t-test. \*\*\**p* = 0.0007. **(G)** V_m_ was not changed by mC4 overexpression. t-test. *p* = 0.2166. **(H)** R_m_ was not affected by mC4 overexpression. t-test. *p* = 0.5455. **(C-H**) Control: N =16 cells; mC4: N = 17 cells. Blue data points: control. Red data points: mC4. Mean ± SEM.

### mC4 overexpression enhances microglia-neuron interactions during development

Several lines of evidence suggest that microglia control circuit maturation and synaptic refinement through the secretion of molecules and direct interactions with neurons (Wu et al., 2015). To test the hypothesis that microglia-neuron interactions contribute to C4-mediated reduction in functional connectivity of L2/3 pyramidal neurons at P21, we combined IUE with histological labeling of microglia. As a proxy for quantifying microglia contact with neurons, we measured the colocalization of neuronal electroporated GFP with a microglia stain (Iba1) in single Z planes using confocal microscopy (Figure 5A). We found instances of GFP puncta colocalized with microglia in both conditions (Figure 5B; white arrowheads), suggesting that microglia were in close contact with the dendritic processes of transfected L2/3 neurons. Neuronal C4 overexpression caused a ~35% increase in the number of microglia that colocalized with GFP puncta and a ~2-fold increase in the area of GFP signal that colocalized with the microglia cell body and proximal processes (Figure 5C and D). These results suggest that increased expression of C4 in cortical neurons enhanced microglia-neuron interactions in dendrites located in superficial L1 and L2/3.

**Figure 5:**
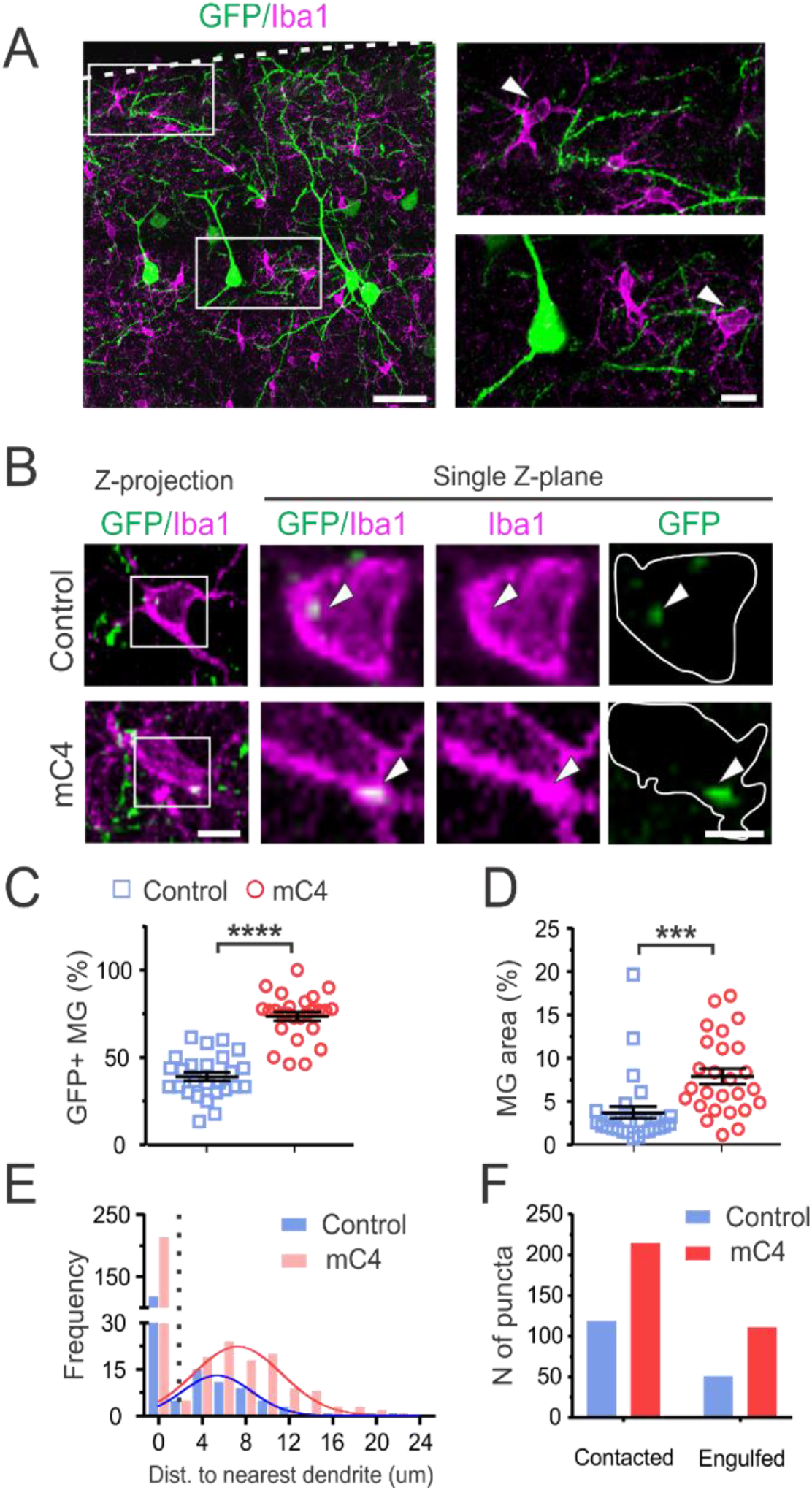
mC4 overexpression enhances microglia-neuron interactions during development. **(A)** Representative single z plane confocal image (60X) showing microglia interacting with processes of electroporated neurons in superficial layers of mPFC (left). White dotted line: pia. Left scale bar = 50 μm. Higher magnification of insets (left) of L1 and L2/3. White arrowhead: microglia (MG). Right scale bar = 10 μm. **(B)** Representative confocal image (60X) showing microglia (Iba1, magenta) colocalized with neuronal GFP signal in P21 histological sections for control (top) and mC4 (bottom) conditions. White arrow heads: Iba1/GFP-positive puncta in microglia soma. Left: max Z-projection of entire microglia; scale bar: 7 μm. Right: single z-plane; scale bar = 3.5 μm. **(C)** Overexpression of mC4 in L2/3 neurons increases the number of microglia colocalized with GFP+ neuronal material (GFP-positive microglia (%)). t-test. \*\*\*\**p* < 0.0001. **(D)** mC4 overexpression increases the percentage microglia area colocalized with neuronal material (MG area (%) = Area of MG GFP+ / Total MG Area). Data represent mean ± SEM. t-test. \*\*\**p* = 0.0009. (E) Histogram plotting the frequency of Iba1/GFP-positive puncta as a function of distance to the nearest dendritic process. Non-zero peaks and values were fitted to Gaussian fits (Control: blue line, peak at 5.38 μm. mC4: red line, peak at 7.27 μm). Dotted line: 1.8 μm (2X optical resolution in the z-axis (0.9 μm). **(F)** mC4 overexpression increased the overall number of Iba1/GFP-positive puncta by ~2 fold, but the proportion of contacted versus engulfed puncta was unaffected. Same puncta as in (D) but sorted into contacted (distance to nearest dendrite = 0 μm) or engulfed (distance to nearest dendrite > 0 μm). Chi-square test. *p* = 0.36. **(C-F)** N = 26 ROIs from 4 mice per condition (including 373 control microglia and 334 mC4 microglia). Mean ± SEM.

The C4-dependent increase in microglia colocalization with GFP could be due to microglia closely interacting with the dendritic processes of transfected neurons; alternatively, colocalization of signals could be the result of microglia engulfment of neuronal material. To further characterize microglia-neuron interactions, we sorted GFP signal colocalized with microglia as either contacted puncta (if microglia were closely interacting with neuronal processes) or as engulfed puncta (if GFP puncta was isolated from neuronal processes) (Figure 5B). To do this, we observed GFP-positive puncta through a Z-stack image to identify “contacted puncta” that were continuous with a neuronal process (distance to nearest neuronal process = 0 μm). If the double-stained puncta were not connected to a neuronal process, they were identified as “engulfed puncta” and the distance from the puncta to the nearest neuronal process was measured along the 3-dimensional space (distance to nearest neuronal process > 0 μm). In control conditions, the average distance from engulfed puncta to the nearest neuronal process was 5.44 μm (obtained from Gaussian fit), which is ~6 times greater than the optical resolution limitation of our confocal microscope for these images (Z optical resolution = 0.9 μm) (Figure 5E). Since the engulfed puncta were isolated from nearby dendrites, we determined that they were not instances of microglia in close contact with neuronal processes. mC4 overexpression led to a ~1.8-fold increase in contacted puncta and a ~2.2-fold increase in engulfed puncta (Figure 5F). However, increased C4 expression did not lead to a change in the proportion of microglia interactions versus microglia engulfment between control and mC4 conditions (Figure 5F). Taken together, our data show that increased levels of neuronal C4 leads to aberrant microglia-neuron interactions and enhanced microglia-mediated engulfment of neuronal material during early postnatal development.

### mC4 overexpression enhances microglia engulfment of postsynaptic PSD-95

Since the gross morphology of L2/3 neurons was not altered by the overexpression of C4 (Supplemental Figure 3), we evaluated whether microglia engulfment of neuronal material (Figure 5) was specifically due to the phagocytosis of synaptic material. To do this, we quantified the colocalization of PSD-95 within lysosomes (CD68), an organelle in microglia that degrades phagocytosed material (Schafer et al., 2012). Endogenous PSD-95 was fluorescently labeled *in vivo* using PSD95-FingR (EF1a-PSD95.FingR-RFP) (Gross et al., 2013) which was introduced to neurons using IUE for both control and mC4 conditions. This labeled only endogenous PSD-95 within transfected neurons with RFP (Figure 6A; pseudocolored green). Fluorescent signal from PSD95-FingR was constrained to superficial L1 and L2/3 including transfected cell bodies of L2/3 neurons, indicating that when this construct is expressed in L2/3 pyramidal neurons, it recapitulates endogenous PSD-95 expression pattern (Hunt et al., 1996; Gray et al., 2006) (Figure 6A and B). In support of this, higher magnification confocal images showed fluorescently labeled puncta in L1 and L2/3 that were not present in L4, which is consistent with the localization of L2/3 dendritic arbors (Figure 6C; white arrowheads).

**Figure 6:**
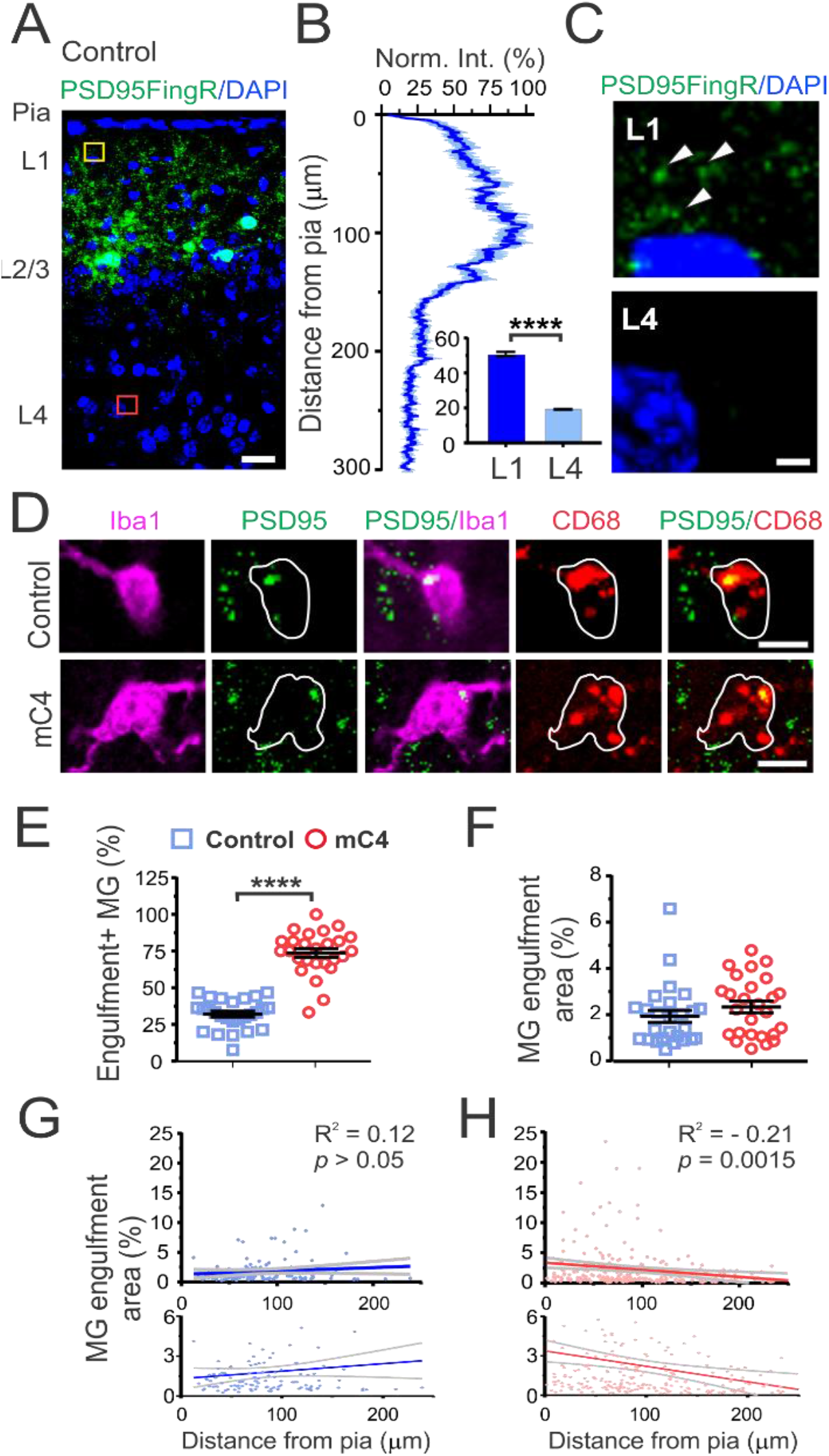
mC4 overexpression enhances microglia engulfment of postsynaptic PSD-95. **A)** Representative confocal image (60X) showing cytoarchitecture (DAPI, blue) and labeling of endogenous PSD-95 by FingR (PSD95FingR, RFP, pseudocolored green) in P21 coronal sections. Scale bar = 25 μm. **(B)** PSD95FingR labeling pattern is consistent with endogenous location of synaptic PSD-95 in L2/3 pyramidal neurons. Mean normalized fluorescent intensity (normalized to peak PSD95FingR signal) as a function of distance from pia (μm). Dark blue line: mean. Light blue shade: SEM. Bar graph: mean normalized fluorescent intensity is greater in L1 (yellow inset) than L4 (red inset). t-test. *p* < 0.0001. **(C)** Zoomed images from image in (A) showing boxed regions in L1 (yellow inset) and L4 (red inset). White arrowheads: PSD95FingR puncta. Blue: DAPI (cell nuclei). Scale bar = 5 μm. **(D)** Representative confocal image (60X) showing PSD-95 located in microglial lysosomes in P21 mice for control (top panels) and mC4 conditions (bottom panels). Magenta: microglia (Iba1). Red: lysosomes (CD68). Green: PSD-95 (PSD95FingR-RFP, pseudocolored green). White line: outline of microglia soma. Single z-plane shown. Scale bar = 10 μm. (E) mC4 overexpression increased the number of microglia positive for PSD-95 engulfment (PSD95FingR signal colocalized with CD68 and Iba1). t-test. \*\*\*\**p* < 0.0001. **(F)** There was no difference in area of PSD95 colocalized with microglia lysosomes between conditions. (MG engulfment area % = area of microglia occupied by PSD95FingR signal in lysosomes / Total microglia area). t-test. *p* = 0.1927. **(G-H)** Microglia engulfment area (%) (from (F)) as a function of cortical depth (μm) for control (G) and mC4 (H). Bottom graphs are same as top but zoomed on the y-axis. Blue line: control mean. Red line: mC4 mean. Gray lines: 95% confidence intervals. Pearson’s correlation. Control: R^2^ = 0.12. *p* > 0.05. mC4: R^2^ = −0.21. \*\**p* < 0.01. **(E-H)** Control: N = 26 ROIs (from 5 mice; 345 microglia). mC4: N = 26 ROIs (from 5 mice; 319 microglia). Mean ± SEM.

Next, we quantified the co-localization of fluorescent signals from PSD-95, Iba1 and CD68 to measure postsynaptic puncta within the lysosomes of microglia in control and C4 conditions (Figure 6D). mC4 overexpression led to a ~2.3-fold increase in the percentage of microglia positive for triple-stained puncta (Figure 6E). However, there was no change in the area of PSD-95 within the lysosomes of microglia that were positive for engulfment (Figure 6F).

Since increased expression of mC4 led to specific spine loss in L1 apical tufts with no changes in basal dendrites, we tested whether microglia engulfment of synaptic material was more prominent near the apical dendrites of cortical neurons. We measured the distance from the center of each microglia soma to the pia to identify its cortical depth and this measurement was then correlated to the area of triple stained puncta colocalized in each microglia. This allowed us to understand if the connectivity deficits in apical tufts were due to altered microglia phagocytosis of synapses in a layer-specific manner. In control conditions, there was no correlation between amount of PSD-95 co-localized with microglia and cortical depth, suggesting that during normal development microglia were not biased towards phagocytosis of PSD-95 in specific layers (Figure 6G). However, we observed a significant correlation in the mC4 condition such that microglia closer to L1, where apical tufts are located, engulfed more PSD-95 than their counterparts in deeper layers (Figure 6H).

To test if complement-dependent enhancement of phagocytosis in L1 was due to an increased density of microglia, we quantified microglia density in L1 (cortical depth: 0 to 120 μm) and L2/3 (cortical depth: >120 to 300 μm) and found no difference between layers in both conditions (Supplemental Figure 4A). There was also no difference in CD68 area within microglia between control and mC4 conditions (Supplemental Figure 4B). In summary, our data support the hypothesis that C4 overexpression drives layer-specific circuit dysfunction through excessive microglia engulfment of postsynaptic material.

### Complement-dependent alterations in frontal cortical circuitry are sufficient to alter maternal-pup social interactions

To determine whether overexpression of mC4 in the frontal cortex is sufficient to cause deficits in early social behaviors, we performed a task to monitor sensorimotor abilities and maternal-pup social interactions (Zhan et al., 2014). For these experiments, we used a modified IUE method (see methods) to target large populations of L2/3 neurons in both hemispheres of the frontal cortex for control (N=15) and mC4 (N=21) mice. Post hoc analysis of brains from these mice showed that most transfected cells were in prefrontal cortical regions, confirming that we were able to target and increase C4 expression in large populations of PFC L2/3 neurons (Supplemental Figure 5).

To test sensorimotor abilities (Maternal interaction 1, MI1 task), we transferred P18 pups to an arena and placed them in a ‘starting corner’ where they were free to explore for 3 min. The arena contained fresh bedding in two neutral corners and nesting material from the animal’s home cage in the corner opposite to the starting corner (Figure 7A). Similar to controls, mC4 mice spent more time exploring the nest corner relative to the neutral corners suggesting that mC4 mice have normal homing behavior (Figure 7B). Although grooming occurrences of mC4 mice were slightly more frequent and ~4.3-fold longer (Supplemental Figure 6), there was no interaction between experimental group and corner preference (Figure 7B). These results indicate that although mC4 mice engaged in more repetitive behavior, they were able to explore the arena and interact with the nest bedding to the same extent as control animals. These results suggest that increased levels of mC4 in the frontal cortex do not impair overall sensory abilities or cause gross motor deficits in P18 pups.

**Figure 7:**
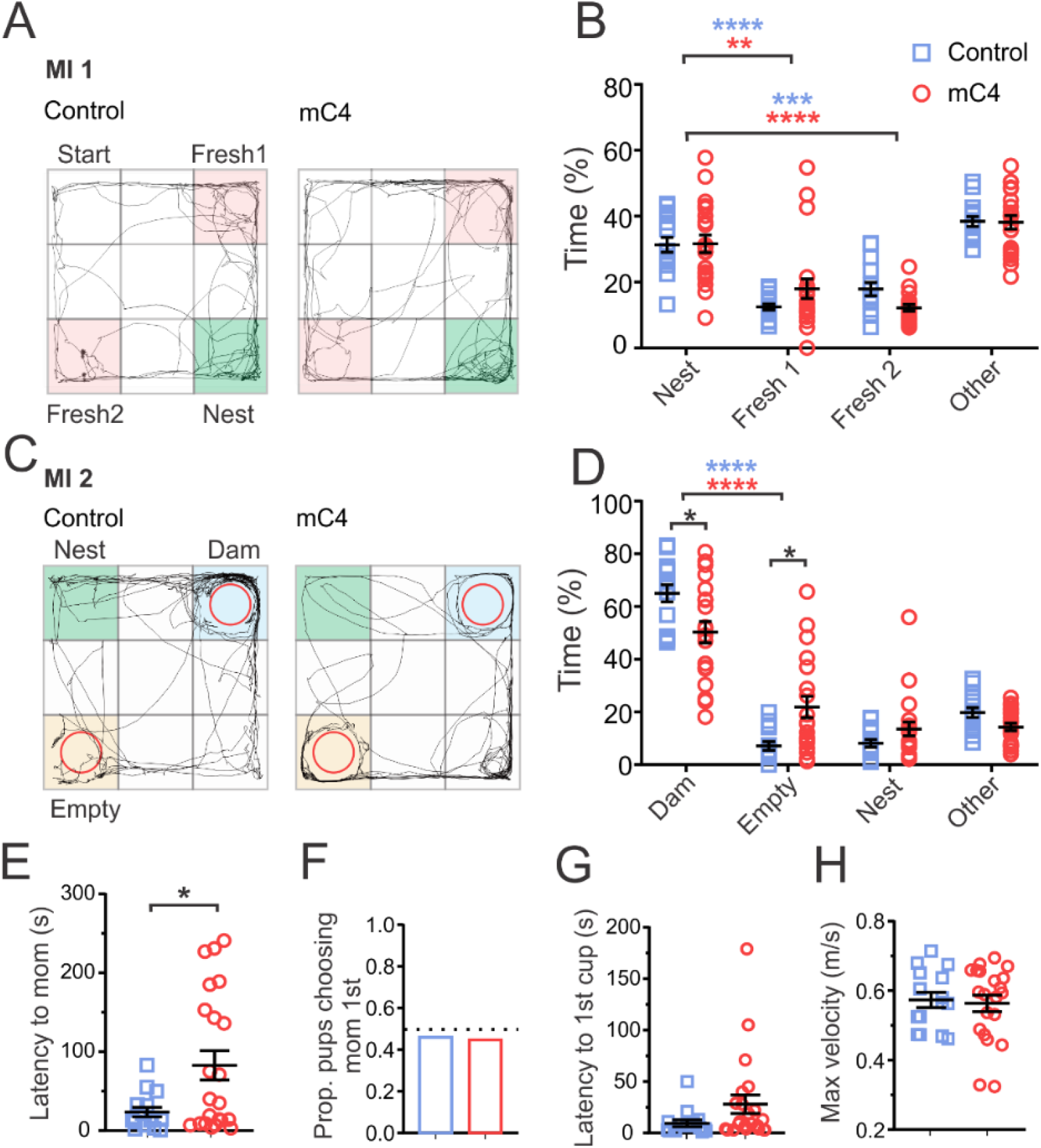
C4-dependent alterations in frontal cortical circuitry are sufficient to alter maternal-pup social interactions. **(A)** Representative examples of path traveled (black trace) by P18 control and mC4 pups in MI1 task. Fresh bedding corners (fresh 1 and 2, pink) and nest bedding corner (green). **(B)** Control and mC4 pups spent a similar proportion of time exploring the nest and fresh corners in the MI 1 task, suggesting motor and sensory skills are intact. All mice spent more time in nest bedding than fresh bedding. \*\**p* < 0.01, \*\*\**p* < 0.001, \*\*\*\**p* < 0.0001. Two-way ANOVA and Sidak’s post test. Control vs. mC4 time spent in nest: *p* > 0.9999. Control vs. mC4 time spent in fresh 1: *p* = 0.3327. Control vs. mC4 time spent in fresh 2: *p* = 0.3138. **(C)** Representative examples of path traveled (black trace) by P18 control and mC4 pups in MI 2 task. Dam’s cup (dam, blue), Empty cup (empty, yellow), Nest bedding corner (nest, green). **(D)** mC4 pups spend less time interacting with dam and more time near the empty cup compared to controls. Two-way ANOVA and Sidak’s post test. \**p* < 0.05. \*\*\*\**p* < 0.0001. **(E)** mC4 mice took longer to approach the dam relative to controls (seconds). t-test with Welch’s correction. \**p* < 0.05. **(F)** Control and mC4 pups traveled to dam’s cup first ~ 50% of the time, suggesting first choice was random. Fisher’s exact test. *p* = 0.9999. **(G)** Latency to reach first cup (either dam or empty) was not different. t-test with Welch’s correction. *p* = 0.06. **(H)** Control and mC4 pups reached the same maximum velocity (m/s). t-test. *p* = 0.77. **(B, D-H)** N=15 control mice and N=21 mC4 mice. Blue: control; red: mC4. Mean ± SEM.

To test maternal-pup social interactions (Maternal interaction 2, MI2 task), we placed P18 pups in an arena that contained nest bedding, an empty wire mesh cup, and a wire mesh cup containing the pup’s dam and the behavior of each pup was monitored for 5 min (Figure 7C). We found that mC4 mice showed a ~23% reduction in time spent exploring the cup containing the dam as compared to control mice (Figure 7D). In agreement with this result, mC4 mice also exhibited a ~2.5-fold decrease in the latency to first approach the dam’s cup (Figure 7E). The proportion of pups that approached the dam’s cup before the empty cup (at start of trial) was close to 50% (Figure 7F), suggesting that pups in both conditions engaged in similar exploratory behavior of the arena before seeking their mother. mC4 and control mice had similar delay times in first reaching either the empty or dam’s cup (Figure 7G) and similar maximum speeds (Figure 7H). Taken together, these results indicate that mC4 pups have intact motor and sensory abilities but are less motivated or interested in interacting with their mother.

## DISCUSSION

The role of the SCZ-associated gene C4 in the development of the mPFC, a brain region that is highly implicated in SCZ pathology, has remained unclear. We show here that overexpression of C4 in the mPFC perturbs cortical circuit development. We demonstrate that, during normal early postnatal development, excitatory neurons in the mouse mPFC express low levels of C4. Moreover, overexpression of C4 in these cells leads to spine dysgenesis that is evident through a transient decrease in filopodia and medium-sized spine density during the third week of postnatal development. We also show that C4-dependent spine abnormalities are accompanied by a decrease in excitatory synaptic drive onto pyramidal cells in the mPFC. C4 overexpression increases microglia-neuron interactions and causes microglia to engulf more PSD-95 of transfected neuron’s apical dendrites in L1, where spine loss is observed. Lastly, increased expression of C4 in the frontal cortex is sufficient to reduce social interactions in juvenile mice. Our findings implicate C4 as playing a role in shaping the developmental trajectory of cortical circuits and provide a causal link between increased C4 levels and prefrontal cortex dysfunction. These data show for the first time that circuit dysfunction in the PFC drives behavioral phenotypes in SCZ rather than merely being a consequence of disease and/or pharmacological treatment.

Our morphological characterization of L2/3 mPFC neurons reveals that C4-mediated connectivity deficits are both circuit and spine-type specific. We show that overexpression of C4 caused a decrease in spine density of apical but not basal dendrites of L2/3 neurons. These data provide evidence that in the mPFC, complement-mediated synaptic alterations are cortical layer and/or input-specific. Interestingly, previous data have shown that exposure to repeated stress in adolescent rats can reduce connectivity in superficial L1 of the mPFC (Negrón-Oyarzo et al., 2015), suggesting that this layer is susceptible to genetic and environmental perturbations during development. When we further investigated which spine-types were affected, we found a specific loss of filopodia and medium-sized spines, while more mature mushroom spines were intact. It is possible that increased expression of C4 does not alter larger, more mature spines perhaps due to their higher levels of synaptic transmission compared to weaker, more immature spines. Large mushroom and stubby spines are also associated with an accumulation of extracellular matrix proteins which could protect them from synaptic elimination by microglia (Levy et al., 2014). Future studies could shed light on the molecular mechanisms of input-specific plasticity by identifying protective signals (Elward and Gasque, 2003; Lehrman et al., 2018) or complement genes that are differentially expressed in large mushroom versus filopodia spine-types. For example, it is possible that CNS activators of C1q, the complement cascade initiator, are enriched in particular brain regions where spine turnover rate is higher (Györffy BA, et al 2018). Although astrocytes and microglia are a source of complement proteins in the brain (Veerhuis et al., 2011), several studies have shown that neurons also express various members of the complement system including C1q, C3 and C4 (Veerhuis et al., 1998, Lévi-Strauss and Mallat, 1987, Gasque et al., 1995, Shen et al., 1997). In this context, our data provide evidence that mPFC neurons are susceptible to increased levels of C4 during development.

Major contributors to synapse formation are dendritic filopodia, thin specialized postsynaptic structures that orchestrate synapse formation by dynamically sampling potential presynaptic partners, thus optimizing circuits (Bonhoeffer and Yuste, 2002; Ozcan, 2017). Given that we see a marked reduction in filopodia and medium-sized spines in C4 overexpressing neurons, it is possible that C4 contributes to aberrant synaptic wiring by reducing the ability of developing circuits to form appropriate connections. The developmental alterations in spine morphology caused by increased levels of C4 could be due to deficits in synapse formation or increased spine elimination, or an interplay between both processes. At its simplest, increased levels of C4 may underlie PFC pathology and dysfunction by specifically altering filopodia-dependent synapse formation during early development. This interpretation suggests that subsequent aberrant synaptic elimination is a consequence of suboptimal circuit wiring. Transient deficits in dendritic spines and filopodia are seen in several mouse models of neurodevelopmental disorders (Meredith, 2015). Similar to these models, our data suggest that there are sensitive developmental periods where cortical circuitry is especially susceptible to altered expression of C4.

Our electrophysiological data strengthens our morphological findings and provides additional insight into the functional abnormalities caused by C4 overexpression. Increased expression of C4 in cortical neurons caused a reduction in the frequency of mEPSCs which parallels developmental dendritic spine dysgenesis. Interestingly, we also observed a decrease in the amplitude of mEPSCs, demonstrating a decrease in the strength of remaining synapses of C4-overexpressing neurons. Previous studies have shown that microglia are able to trogocytose, or “nibble” and phagocytose, small volumes of neurons and synapses (Weinhard et al., 2018). It is possible that microglia trogocytose some of the postsynaptic density of synapses without fully eliminating them, thus causing a reduction in synaptic strength due to a loss of glutamate receptors. In our model, the C4-dependent decrease in excitatory synaptic drive could be the result of an increase in phagocytosis of synapses by microglia. Alternatively, microglia trogocytosis of synaptic material could trigger synaptic elimination mediated through conventional synaptic mechanisms such as long-term depression (Sheng & Erturk, 2013). Although C4 did not induce a change in the frequency or amplitude of mIPSCs, it significantly slowed the kinetics of inhibitory transmission. Previous studies have shown that inhibitory transmission is distributed non-uniformly along dendritic shafts with a higher density of inhibitory synapses onto spines at apical distal dendrites as compared to proximal locations (Chen et al., 2012), suggesting that inhibitory transmission could be regulated by excitatory synapses. During early development, the C4-dependent loss of dendritic spines could alter inhibitory synapse location and thus lead to slower inhibitory currents (Gambino and Holtmaat, 2012). Together with our morphological data, these results support the hypothesis that abnormal levels of C4 cause developmental delay and primarily affect the functional connectivity of neurons by reducing excitatory synaptic drive.

In addition to being the brain’s resident macrophage, microglia are known to influence brain development by playing roles in synapse formation, elimination and maintenance (Wu et al., 2015). Here we show that abnormal levels of C4 during development lead to aberrant microglia-neuron interactions and engulfment of synaptic material. Since microglia are the only cells in the brain that express CR3 (Schafer and Stevens, 2015), they are equipped to recognize material that has been tagged by the complement cascade to initiate phagocytosis. Our data in the PFC are consistent with and expand upon previous studies in the retinogeniculate pathway that show that microglia engulf synapses in a complement-dependent manner (Schafer et al., 2012); suggesting that complement-dependent circuit wiring is necessary for normal development and contributes to pathology when mis-regulated. Complement-dependent alterations in the plasma membrane have been associated with electrophysiological changes in cultured cells (Stephens and Henkart, 1979). Since we did not observe changes in gross morphology in neurons overexpressing C4, the decrease in capacitance we measured could be due to a change in membrane tension or composition due to complement activity or microglia-neuron interactions (Dai et al., 1998). A recent study showed that complement-based synaptic pruning is linked to local apoptotic-like processes at the synapse (Györffy et al 2018), thus supporting the hypothesis that complement pathway activation could alter the properties of the plasma membrane. We propose that neuronal overexpression of C4 contributes to SCZ pathology by causing an excessive number of synapses tagged by the complement cascade to be phagocytosed by microglia, thus leading to reduced connectivity.

Individuals with SCZ exhibit decreased blood flow and metabolism in the frontal cortex during rest and while performing cognitive tasks, known as hypofrontality (Wolkin et al., 1992; Andreasen et al., 1997; Hill et al., 2004). Moreover, the extent of hypofrontality correlates with gray matter loss and SCZ symptom severity (Collin et al., 2012; Vita et al., 2012). Our results show that C4 overexpression leads to a reduction in excitatory synaptic drive in L2/3 cortical neurons, thus resembling a hypofrontality phenotype. Social deficits in individuals with SCZ are an early feature of this disease and poorer social functioning is associated with worse functional outcome in adults (Velthorst et al., 2017; Green, 2016). Investigating the causes for early social deficits could thus reveal potential targets for early therapeutic intervention in schizophrenia. In L1, the apical dendrites of pyramidal cells in multiple cortical layers interact with projections from higher order areas (Douglas and Martin, 2007). It is thought that cortical ‘feedback’ that enters L1 exerts ‘top-down’ control that is important for higher-order cognitive processes (Sjöström and Häusser, 2006); therefore, the increased vulnerability of this layer to C4 overexpression could provide a cellular substrate for social dysfunction. Our data suggest that this loss of structural and functional connectivity contributes to early social deficits in mice. In this regard, the changes we observed in connectivity and behavior parallel known SCZ phenotypes (Glantz and Lewis, 2000; Thompson et al., 2001; Vita et al., 2012; Green, et al., 2015; Velthorst et al., 2017) and provide evidence that the PFC regulates early social behavior.

Although polygenetic mechanisms underlie SCZ (Schizophrenia Working Group of the Psychiatric Genomics Consortium, 2014), we provide evidence that altering expression of a single gene, C4, in mPFC neurons is sufficient to cause known cellular and behavioral phenotypes that resemble SCZ pathology. In summary, we have identified a critical developmental window where prefrontal cortical circuits are susceptible to alterations in C4 expression, thus opening the possibility for early therapeutic intervention to alter the developmental trajectory of SCZ.

## Acknowledgments

We thank Andrzej Cwetsch for input in the design of the tripolar electrode, Todd Blute and the Boston University Biology Imaging Core for use of the confocal microscope, Tim Gardner for use of the cryostat and Byron Price for helpful consultation with behavior analysis. We thank Thomas Gilmore, Ben Scott, Carlos Portera-Cailliau, and members of the Cruz-Martín lab for critical reading of the manuscript and helpful discussions. We used Boston University’s EPIC facilities for 3D printing. This work was supported by a NARSAD Young Investigator Grant (A.C.M.), the Brenton R. Lutz Award (A.L.C.), the NSF NRT UtB: Neurophotonics National Research Fellowship (A.L.C. and L.K.), and the Boston University Undergraduate Research Opportunities (J.L., F.S.H., T.N., K.L.K., E.N., W.W.Y).

## Author Contributions

A.L.C., T.J. and A.C.M. designed experiments. A.L.C. and A.C.M. wrote the paper. A.L.C, T.P.H.N. and L.N.K. performed surgeries. A.L.C. and L.N.K. performed and analyzed *in situ* hybridization experiments. A.L.C., J.L., F.S.H., T.P.H.N. and T.J. performed confocal imaging. A.L.C., T.P.H.N. and J.L. quantified and analyzed spine data. T.J. performed electrophysiological recordings and analyzed data. A.L.C. performed microglia assays and analyzed data. A.L.C., T.J., E.R.N., and L.N.K. performed behavioral experiments. T.J., E.R.N., W.W.Y., S.R. and A.C.M. analyzed behavioral data. A.L.C., F.S.H., K.L.K. and S.R. performed cell quantification and analysis of behavior brains. A.C.M. supervised the project.

## METHODS

### Experimental model details

All experimental protocols were conducted according to the National Institutes of Health (NIH) guidelines for animal research and were approved by the Boston University Institutional Animal Care and Use Committee (IACUC). All mice were group housed on a 12-h light and dark cycle with the lights on a 7 AM and off at 7 PM with food and water *ad libitum*. Offspring were housed with dams until weaning at P21. For spine quantification and microglia interactions and engulfment, electrophysiology and behavior experiments, wild-type CD-1 mice of both sexes from age 7-60 days were used (CD-1 IGS; Charles River; strain code: 022). For *in situ* hybridization, *C4B* knockout mice in a C57BL/6 genetic background (The Jackson Laboratory, stock number: 003643) and wild-type C57BL/6 mice of both sexes at age P30 were used (Charles River; strain code: 027). Males and females were used for all experiments, and no differences were found between sexes.

### DNA constructs

For control conditions we used a plasmid containing EGFP under the CAG promoter (pCAG-EGFP, Addgene plasmid #11150) (Matsuda and Cepko, 2004). DNA sequences containing mouse *C4B* (NM_009780.2, synthesized by Genescript) and human *C4A* (RC235329, Origene) were subcloned (InFusion Kit, Clonetech) into the pCAG backbone to produce pCAG-mC4 and pCAG-hC4A, respectively. Endogenous PSD-95 was fluorescently labeled *in vivo* using PSD95-FingR (EF1a-PSD95.FingR-RFP) (Gross et al., 2013). All DNA was purified using the ZymoPureII (Zymo Research) plasmid preparation kit and resuspended in molecular biology grade water.

### Antibodies and histology

Mice were administered a lethal dose of sodium pentobarbital (250 mg/kg; IP) before being transcardially perfused with PBS and then 4% PFA solution. After brains were dissected and post-fixed for 24 h in 4% PFA solution, they were transferred to a 30% (w/v) sucrose solution and stored at 4° C. Coronal sections were cut on a sliding microtome (Lecia SM2000) at different thicknesses appropriate for each experiment (see specific methods). For immunostaining, tissue was blocked and permeabilized in 10% donkey serum with 0.25% TritonX100. Sections were incubated with primary antibodies overnight at 4°C on a shaker, and with secondary antibodies for 2 h at room temperature. After incubation periods and between each step, sections were rinsed three times with PBS for 10 min. Depending on the experiment, the following primary antibodies were used: guinea pig anti-Iba1 (Synaptic Systems, 234004, 1:500), and rabbit anti-CD68 (Abcam, ab125212, 1:500). The following secondary antibodies were used: donkey anti-guinea pig Alexa Fluor 594 (Jackson Laboratories, 706-545-148, 1:1000), donkey anti-guinea pig Alexa Fluor 488 (Jackson Laboratories, 706-585-148, 1:1000), and donkey anti-rabbit Alexa Fluor 647 (Jackson Laboratories, 711-605-152, 1:1000). Brain sections were subsequently mounted onto microscope slides (Globe Scientific) using Fluoromount-G mounting medium with DAPI (Thermo Fisher Scientific).

### *In utero* electroporation

L2/3 progenitor cells in the mPFC were transfected via *in utero* electroporation (Szczurkowska et al., 2016). pCAG-EGFP was electroporated in the control condition, and pCAG-EGFP plasmid was co-electroporated with either pCAG-mC4 or pCAG-hC4A plasmids for the mC4 or hC4 groups, respectively. Prior to surgery, all tools were sterilized by autoclaving. Aseptic techniques were maintained throughout the procedure and a sterile field was prepared prior to surgery using sterile cloth drapes. Animals were weighed and a combination of buprenorphine (3.25 mg/kg; SC) and meloxicam (1-5 mg/kg; SC) were administered as preoperative analgesics. Timed-pregnant female CD-1 mice at embryonic day 16 (E16) were anesthetized by inhalation of 4% isoflurane and maintained with 1-1.5% isoflurane via mask inhalation. The abdomen was sterilized with 10% povidone-iodine and 70% isopropyl alcohol (repeated 3 times) before a vertical incision was made in the skin and then in the abdominal wall. The uterine horn was then exposed to allow injection of 0.5-1.0 μl of DNA solution (containing 1 μg/μl plasmid and 0.1% Fast Green) into the lateral ventricles using a pressure-injector (Picospritzer III, Parker Hannifin) with pulled-glass pipettes (Sutter Instrument, BF150-117-10). To target L2/3 progenitor cells in prefrontal cortex for imaging and electrophysiological experiments, a custom-built triple electrode probe (Szczurkowska et al., 2016) was placed by the head of the embryo, with the negative electrodes placed near the lateral ventricles and the positive electrode placed just rostral of the developing prefrontal cortex. For bilateral *in utero* electroporations used in behavioral experiments, DNA was injected into both lateral ventricles by positioning the glass pipette at a 90-degree angle relative to the midline of the embryo’s head and injecting 2-4 μl of DNA solution. These modifications ensured that the DNA solution would travel to the lateral ventricle contralateral to the injection site. Next, 4 square pulses (pulse duration: 50 ms, pulse amplitude: 36 V, inter-pulse interval: 500 ms) were delivered to the head of the embryo using a custom-built electroporator (Bullmann et al., 2015). Embryos were regularly moistened with warmed sterile PBS during the surgical procedure. After electroporation, the embryos and uterine horn were gently placed back in the dam’s abdominal cavity and the muscle and skin were sutured (using absorbable and non-absorbable sutures, respectively). Finally, the dams were allowed to recover in a warm chamber for 1 h and then returned to their cage.

### Imaging

Fluorescence images were collected using an inverted laser scanning confocal microscope (Nikon Instruments, Nikon Eclipse Ti with C2Si+ confocal) controlled by NisElements (Nikon Instruments, 4.51) including four laser lines (405, 488, 561, and 640 nm). For M-FISH and microglia experiments, confocal images were taken with a 60X Plan Apo objective (Nikon Instruments; Plan Apo, NA 1.4, WD: 0.14 mm, oil objective) using 1024 x 1024 pixel scans (pixel size = 0.27 x 0.27 μm). For dendritic spine imaging, images from P7-9, P14-16, P21-23 and P55-60 brain tissue were taken using a 40X Plan Apo λs objective (Nikon Instruments; Plan Apo, NA 1.3, WD: 0.2 mm, water objective) using 1024 x 1024 pixel scans (pixel resolution = 0.12). We imaged apical dendritic tufts in L1 and basal dendrites in L2/3. For dendritic spine imaging and neuronal reconstructions, we collected ROIs that consisted of stacks of images (~20-40 optical sections, z-step = 0.3 μm and 1 μm, respectively). The area and diameter of neurons were analyzed in 40X images from the brightest z-plane of the soma. Brain sections from behavior mice were imaged using an upright wide-field microscope (Nikon Instruments, Nikon Eclipse Ni) controlled by NisElements (Nikon Instruments, 4.20) using a 10X objective (NA: 0.3, WD: 16 mm). DAPI and GFP-transfected cells in behavior brains were imaged using fluorescence filters for BFP (excitation: 370-401, dichroic: 420) and GFP HC (excitation: 470/40, dichroic: 495). In some of the figure panels for dendritic spine images, distracting processes (e.g., axons) were digitally removed in ImageJ (NIH) for display purposes only. Images in all conditions were collected using similar imaging conditions and exposure settings. Imaged stacks were imported in TIFF format into ImageJ (NIH). For confocal image analysis, we only measured neurons and glia in the anterior cingulate cortex, prelimbic, infralimbic and medial orbital divisions of medial prefrontal cortex. For behavior brain analysis, all GFP-positive cells were counted to quantify the extent of electroporation.

### Multiplex fluorescence in situ hybridization

For multiplex fluorescence *in situ* hybridization (M-FISH) experiments, P30 brains were collected and immediately fresh frozen on dry ice in O.C.T. compound (Fisher HealthCare, 23-730-571). Tissue was cut on a cryostat (Leica CM 1800) at 15 μm thickness, and M-FISH experiments were completed by using a commercial assay (RNAscope, Advanced Cell Diagnostics. Inc.). *In situ* fluorescent probes were used to detect mC4 (#445161-C1), EGFP (#400281-C2) and CaMKIIα (#445231-C3). M-FISH assay and all reagents were obtained from Advanced Cell Diagnostics. All M-FISH experiments had N = 3 per condition. For *in situ* analysis, we quantified fluorescent signal from L2/3 CaMKIIα+ cell bodies of transfected GFP+ cells and their untransfected neighbors in mC4 conditions from single z planes of 60X confocal images. This approach allowed us to control for variability of transcript expression between mice. We use the DAPI signal to identify the cell’s nucleus and all quantification of *in situ* signals was performed in the soma’s brightest focal plane. CaMKIIα signal was used to delineate the perimeter of the cell body of L2/3 excitatory neurons in the mPFC. Transfected cells were identified by presence of GFP mRNA, and non-transfected cells were GFP(-). Next, fluorescent signal from GFP and C4 transcripts were thresholded and binarized. The binarized signal was then used to calculate the percentage of soma area covered by GFP or C4 mRNA. We used the same binarization threshold and analysis procedure for both conditions.

### Dendritic spine analysis

For the dendritic spine developmental time course experiments in control, mC4 and hC4 conditions, we analyzed dendritic spines at multiple postnatal days (P7-9, P14-16, P21-23, P55-60). For all groups, we analyzed 10 to 12 dendrites from 3 to 5 mice (from multiple litters). Confocal image z-stacks (40X) were background subtracted and median filtered (radius 0.25 μm), and the presence or absence of a protrusion was determined by visually inspecting the entire z-stack of images. For a dendritic protrusion to be counted, it had to clearly protrude out of the shaft by at least 3 pixels (~0.36 μm). Dendritic protrusions were quantified from either apical dendritic tufts in L1 or secondary/tertiary basal dendrites in L2/3. Spine density was calculated by dividing the number of counted spines by the total dendritic length analyzed (~50-80 μm long shafts). Since GFP brightness is monotonically related to the volume in each protrusion (Holtmaat et al., 2005), we quantified the fluorescent intensity (total integrated brightness, TIB (a.u.)) of dendritic spines in apical tufts. To obtain a TIB value, the mean gray value of the dendritic spine was measured at the brightest focal plane and was divided by the mean fluorescence intensity of the adjacent dendritic shaft to normalize for varying imaging conditions. We sorted dendritic spine types into large mushroom/stubby (TIB > 75%), medium size spines (25% ≤ TIB ≤ 75%) and thin spine/filopodia (TIB < 25%) intensity groups based off of percentile cutoffs determined from TIB distribution values of dendritic spines. It has been previously shown that this is a reliable unbiased approach for classifying dendritic spine types (Cruz-Martín et al. 2010).

### Electrophysiological recordings and analysis

Mice (P18-25) were anesthetized with 4% isoflurane-oxygen mixture (v/v) and perfused intracardially with ice-cold external solution containing the following (in mM): 73 sucrose, 83 NaCl, 26.2 NaHCO_3_, 1 NaH_2_PO_4_, 22 glucose, 2.5 KCl, 3.3 MgSO_4_, 0.5 CaCl_2_ and were bubbled with 95% O_2_/5% CO_2_ (295-305 mOsm). Coronal slices (300 μm thickness) were cut on a VS1200 vibratome (Leica) in ice-cold external solution, before being transferred to ACSF containing the following (in mM): 119 NaCl, 26 NaHCO_3_, 1.3 NaH_2_PO_4_, 20 glucose, 2.5 KCl, 2.5 CaCl_2_, 1.3 MgCl_2_, bubbled with 95% O_2_/5% CO_2_ (295-305 mOsm). Slices were kept at 35 °C for 30 min, before being allowed to recover for 30 min at room temperature. All recordings were performed at 30-32 °C. We only recorded from transfected neurons in the anterior cingulate cortex, prelimbic, infralimbic and medial orbital divisions of the medial prefrontal cortex. Signals were recorded with a 5X gain, low-pass filtered at 6 kHz and digitized at 10 kHz using a patch-clamp amplifier (Molecular Devices, Multiclamp 700B).

Whole-cell voltage clamp recordings were made using 3-5 MΩ pipettes filled with an internal solution that contained 125 mM Cs-gluconate, 3 mM NaCl, 8 mM CsCl, 4 mM EGTA, 4mM MgATP, 0.3 mM NaGTP and 10 mM HEPES, pH 7.3 with CsOH (280-290 mOsm). Series resistance (R_s_) and input resistance (R_in_) were monitored throughout the experiment by measuring the capacitive transient and steady-state deflection in response to a −5 mV test pulse, respectively. For mEPSC recordings, cells were voltage clamped at E_rev_ GABA_A_ (−70 mV) in the presence of 1 uM Tetrodotoxin (Tocris). For mIPSC recordings, cells were voltage clamped at E_rev_ Glu (+5 mV) in the presence of 1 uM Tetrodotoxin (Tocris). For all groups, we analyzed 10 to 12 cells from 4 mice (from multiple litters). mPSCs were detected by fitting to mPSC amplitude template using pClamp10 analysis software (Molecular Devices). The peak current of each mPSC was calculated. Next, average mPSC amplitude and inter-event-interval was calculated for each cell and then averaged across each condition to determine the population mean and SEM. Bath application of 100 uM GABA_A_ inhibitor picrotoxin at E_rev_ Glu and 20 mM CNQX and 50 mM DL-AP5 at E_rev_ GABA_A_ eliminated mPSCs, thus confirming the identity of the recorded currents. Cells were excluded if Rs varied by more than 20% during a recording. For all recordings, series resistance was close to 10 MΩ (control, 9.39 ± 0.53 MΩ, n = 12, mC4, 9.74 ± 0.44 MΩ, n = 10, *p* > 0.05) and was not compensated.

In current-clamp recordings, CNQX (20mM), 50 mM DL-AP5 and picrotoxin (100 μm) were routinely added to the extracellular solution to block ionotropic synaptic transmission mediated by glutamate and GABAA receptors, respectively, to assess persistent firing. Neuronal excitability was assessed using the input–output curve measured from the changes in membrane potential (presence of APs) evoked by current steps (from Vrest, start = −200 pA, step duration: 300 ms), increasing in increments of 10-15 pA in current-clamp mode. Cm and Rm were calculated in seal test configuration from the decay and steady state current of a transient generated in response to a – 5 mV pulse test. Rheobase is the minimum current amplitude (300 ms) that resulted in an AP.

### Neuronal morphology analysis

Confocal images of L2/3 GFP-positive neurons in mPFC (P21 and P60) were collected using a 40X objective. Z-stacks for large field of view images were taken at z-step of 1 μm through the entire section (~200 μm thickness), taking care to include the entirety of the dendritic tree. Image stacks of dendritic trees and soma were imported in TIFF format into ImageJ (NIH). Cell somas and dendrites were traced manually to ensure accurate reconstruction. Dendritic arbors that could not be confidently and completely reconstructed were not used in the analysis. Dendritic reconstructions were confirmed by comparing independent reconstructions from two investigators (N = 10 neurons per condition). The dendritic parameters analyzed included the total dendritic length (of all dendrite branches), number of dendritic branches, number of branch points, number of dendritic end tips. Branch order for each dendritic branch was assigned starting at the cell body and increased after each branch point and the maximum branch order quantified. Sholl analysis was performed using Simple Neurite Tracer (SNT, ImageJ plugin, Longair MH, 2011). Each reconstructed neuron was thresholded and binarized and the skeletons were imported to SNT. Sholl analysis was performed by calculating the number of dendrites that intersected concentric spheres that radiated from the soma in 10-μm radius increments. On some occasions, the dendritic arbor could not be reconstructed for several reasons: dendritic processes extended outside of the acquired field of view (or Z-stack), the field of view became too dense with GFP-positive processes from neighboring cells, preventing unambiguous reconstruction, and some dendritic processes were deemed insufficiently bright to allow for unequivocal reconstruction. Thus, our reconstructions are an underestimate of the entire dendritic arbor. Neuronal soma area and diameter were measured from a single Z-plane for each neuron. Soma diameter was measured perpendicular to the apical axis. Analysis was performed at P21 in the mPFC and 315 neurons were analyzed for control (6 ROIs from 3 mice) and 216 cells were analyzed for the mC4 condition (7 ROIs from 3 mice).

### Microglia assays

#### Colocalization of GFP and Iba1

GFP colocalization with microglia (Iba1) was quantified at P21. Confocal images of mPFC (60X objective) were collected using the same exposure settings between conditions including channels for DAPI, GFP (transfected neurons) and Iba1 (microglia). All images were background subtracted and binarized using the same settings between experimental groups. Analysis was completed for single Z-planes. To identify microglia, ROIs were drawn around the perimeter of the microglia, including the cell body and any proximal processes in the single analyzed Z-plane. Microglia cell bodies were confirmed by the presence of DAPI. Colocalized GFP and Iba1 signal was isolated and quantified within the microglia by creating a thresholded mask. Colocalized signal within the microglia ROIs were only included in the analysis if they were equal to or larger than 1.23 μm (~3 pixels). GFP-positive puncta within microglia were visually inspected in three-dimensional space by examining multiple Z-planes to confirm the containment of the puncta to the microglia cytoplasm. The percentage of colocalized signal was quantified by dividing the area of the mask by the total area of the microglia ROI (GFP/Iba1 colocalization (%) = microglia area colocalized with GFP / total microglia area). Using this analysis, we also calculated the percentage of microglia that were positive for GFP (GFP-positive microglia (%) = total number of GFP-positive microglia / total number of microglia). N = 26 ROIs from 4 mice per condition (including 373 control microglia and 334 mC4 microglia).

#### Nearest neuronal processes analysis

The distance from each GFP-positive puncta colocalized within microglia to the nearest neuronal process was measured by examining Z-stack images (50 μm, z step = 0.5 μm). Double-stained puncta that were continuous with a dendritic process were identified as attached (distance = 0 μm). If the double-stained puncta were not connected to a neuronal process, we then measured the distance from the puncta to the nearest neuronal process along the 3-dimensional space (not attached, distance > 0 μm). N = 26 ROIs from 4 mice per condition (including 373 control microglia and 334 mC4 microglia).

#### Synaptic engulfment and cortical depth correlation

To quantify engulfment of synaptic material, we electroporated (E16) a plasmid that labels endogenous PSD-95 (EF1a-PSD95.FingR-RFP) for control and mC4 conditions. Analysis was completed for single Z-planes using DAPI, PSD-95-FingR-RFP, Iba1 and CD68 (a lysosomal marker). Analysis was completed using the same method as in the GFP and Iba1 colocalization analysis except synaptic material was only counted if it was within the lysosome of the microglia (PSD-95-postive puncta that colocalized with both Iba1 and CD68 signals). For each thresholded mask, the total colocalized area (microglia engulfment area (%) = microglia area colocalized with PSD-95 and CD68 / total microglia area) was quantified. We also quantified the percentage of microglia positive for engulfment (engulfment-positive microglia = (%) total number of positive microglia / total number of microglia). Lastly, the percent area of each microglia colocalized with CD68 signal was calculated as a measure of microglia reactivity. In a separate analysis, the cortical depth of each engulfment-positive microglia was determined by measuring the distance from the center of the cell body (using DAPI) to the adjacent pia mater. We restricted cortical depth analysis to layers 1 and 2/3 of the mPFC, where the dendritic spines of L2/3 cortical neurons are localized (depth ≤ 300 μm). Microglia that did not contain triple-stained puncta (PSD-95/Iba1/CD68) were excluded from correlation analysis. Control: N = 26 ROIs (from 5 mice; 345 microglia). mC4: N = 26 ROIs (from 5 mice; 319 microglia).

#### Microglia density

Microglia density analysis (P21) was completed using 50 μm z-stack images using DAPI and Iba1 signals. Microglia density was independently calculated for L1 (cortical depth: 0 to 120 μm) and 2/3 (cortical depth: >120 to 300 μm) of the mPFC. Density was calculated per ROI by dividing the number of microglia in the ROI by the total volume of tissue for each ROI. Microglia density was measured in 19 control ROIs (from 5 mice; 2,146 microglia) and 17 mC4 ROIs (from 5 mice, 1640 microglia) in the superficial layers of cortex and in each ROI the density was calculated for L1 and L2/3. Microglia cell bodies were confirmed by the presence of DAPI.

### Maternal interaction task

For behavior tasks, we used control (N = 15) and mC4 (N = 21) mice that were electroporated bilaterally at E16. Prior to the maternal homing test, dams were acclimated to the behavioral task by placing them in the mesh cup for 5-min periods, for three consecutive days. Mice were separated from their dams for an hour immediately before testing. The maternal interaction (MI) task consisted of two phases. In phase 1 (MI1) mice were placed in an open field with home bedding and fresh bedding, and in phase 2 (MI2) mice were placed in an open field with home bedding, and their dam restrained in a small wire mesh cup and an empty wire mesh cup. Both phases of the behavioral task were run on P18.

For MI1, individual pups were transferred to a homemade acrylic arena (50 × 50 × 30 cm length-width-height) and placed in a ‘starting corner’. The arena contained fresh bedding in two neutral corners and nest bedding in the corner opposite from the starting corner. Mice explored for 3 min and the total time spent in the starting, nest and fresh corners was measured using a homemade video tracking system written in MATLAB (MathWorks). Grooming occurrences and time spent grooming were annotated by a trained experimenter blind to experimental conditions.

For MI2, two wire mesh cups were placed in opposite corners of a homemade acrylic arena (50 × 50 × 30 cm length-width-height), one containing the animal’s dam and the other empty, and the start corner contained soiled home bedding. Mice spent 5 minutes exploring the MI2 environment and we video recorded behavior using a Logitech C270 Webcam at 30 fps. Time spent near each cup and other areas was measured using our homemade video tracking system. Behaviors were recorded under a dim light (~20 lux) positioned over the center zone of the arena. Between trials and mice, the arena was cleaned with 70% ethanol.

### Quantification of behavior brains

We counted the number of GFP-positive cells and assessed the extent of cell transfection in brains of animals that were tested in the maternal interaction task. Experimental groups included mice from at least two litters (N=15 mice for control and N=21 for mC4 condition).

Somas were counted by two trained independent experimenters and we did not find differences in their quantification. Coronal sections (50 μm thickness) from the entire brain were carefully inspected and the DAPI signal was used to corroborate the presence of cell nuclei. We used a brain atlas (Paxinos and Franklin, 2001) to confirm the location of GFP+ cell bodies and to delineate the boundaries of each brain region. Most transfected cells were located in frontal cortical regions, between Bregma +3.0 mm to 1.6 mm (Figure S1). We did not observe a significant number of cells in caudal cortical regions or subcortical areas.

To estimate the total amount of cells transfected in each brain, cells were counted from 50 μm sections including every other section. Since transfections were homogenous across neighboring brain sections, once we obtained the total number of cells per section, we plotted cell counts against their respective Bregma coordinates. We then used a linear interpolation to estimate the cell counts from missing brain sections. In a small number of brains (N=3), we found no differences in total cell counts when we compared values obtained from counting GFP-positive cells in all transfected brain sections to estimated values obtained using the interpolation approach. All control and mC4 brains from mice used in behavioral tasks were GFP+ and had similar rostro-caudal patterns of transfected neurons. We did not find any differences in total cell counts between groups (Figure S2, control, 3920 ± 678.3 cells, mC4, 4761 ± 812.6 cells, t-test, *p* = 0.439).

### Statistical analysis

For confocal image analysis and electrophysiological recordings, we focused on neurons in the anterior cingulate cortex, prelimbic, infralimbic and medial orbital divisions of medial prefrontal cortex. All statistical analysis was completed in GraphPad Prism 8.0 and threshold for significance for all tests was set to 0.05 (α = 0.05). M-FISH data sets were analyzed with an unpaired t-test. Spine developmental data was analyzed using a two-way ANOVA followed by Tukey’s test. Dendritic spine fluorescent intensity values were sorted based off of percentile cutoffs determined from TIB distribution values of dendritic spines in control conditions, and differences in the density of spine-types between groups were tested with a one-way ANOVA followed by Tukey’s post-test. Electrophysiological data were analyzed with an unpaired t-test and cumulative distributions were analyzed with a Kolmogorov-Smirnov (KS) test. Percent of microglia positive for engulfment and percent of microglia area positive for engulfment were analyzed using an unpaired t-test. Microglia puncta analysis for attached and unattached assay was analyzed using Chi-square test. Behavioral data were analyzed with a two-way ANOVA followed by Sidak’s post-test and a t-test with Welch’s correction. Dendrite and soma morphology measurements were analyzed using a t-test. Both male and female mice were used and we did not observe any differences between the sexes among groups. Correlations were determined using Pearson’s R correlation and linear regression. Analysis was performed blind to condition. Soma area/diameter and dendrite measurements were analyzed using GraphPad Prism 8.0 (GraphPad Software Inc.). Figures were prepared using CorelDRAW Graphics Suite X8 (Corel Corporation) and ImageJ (NIH). Custom written routines for behavioral tracking and analysis are available upon request. Data are presented as the mean ± SEM, unless otherwise noted.

### Data and Software Availability

Custom written routines for behavioral tracking and analysis are available upon request. The datasets generated during and/or analyzed during the current study and all custom scripts and functions generated or used during the current study are available from the lead contact on request.

