## Supplemental Figures for "Increased expression of schizophrenia-associated gene C4 leads to hypoconnectivity of prefrontal cortex and reduced social interaction"

**Supplemental Figure 1**

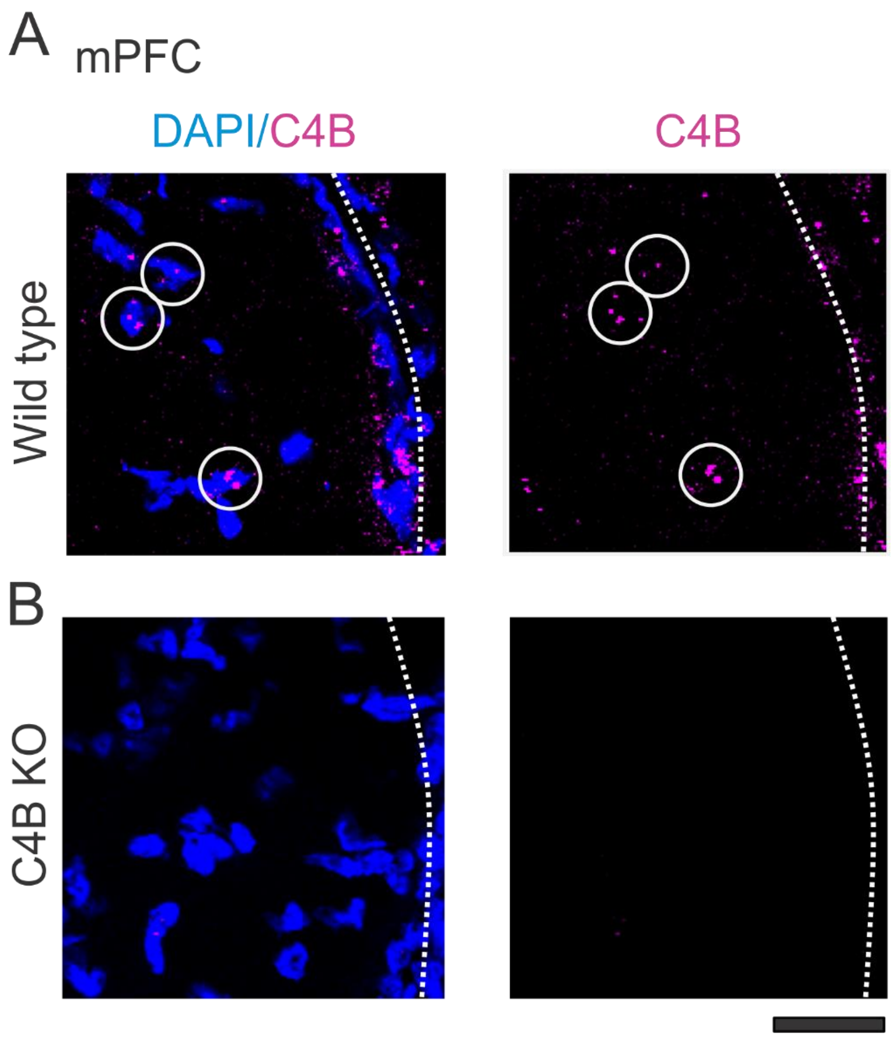


**Supplemental Figure 1: The *in situ* hybridization probe for mC4 is specific.**
**(A)** C4B mRNA is expressed in mPFC superficial layers in P30 WT mice. White dotted line: pia mater. White circles: nuclei with C4 mRNA. Scale bar = 60 µm. **(B)** C4B mRNA is not expressed in mPFC superficial layers in P30 C4B KO mice. White dotted line: pia mater. Scale bar = 60 µm. **(A-B)** Confirmed in 3 mice per condition.

**Supplemental Figure 2**

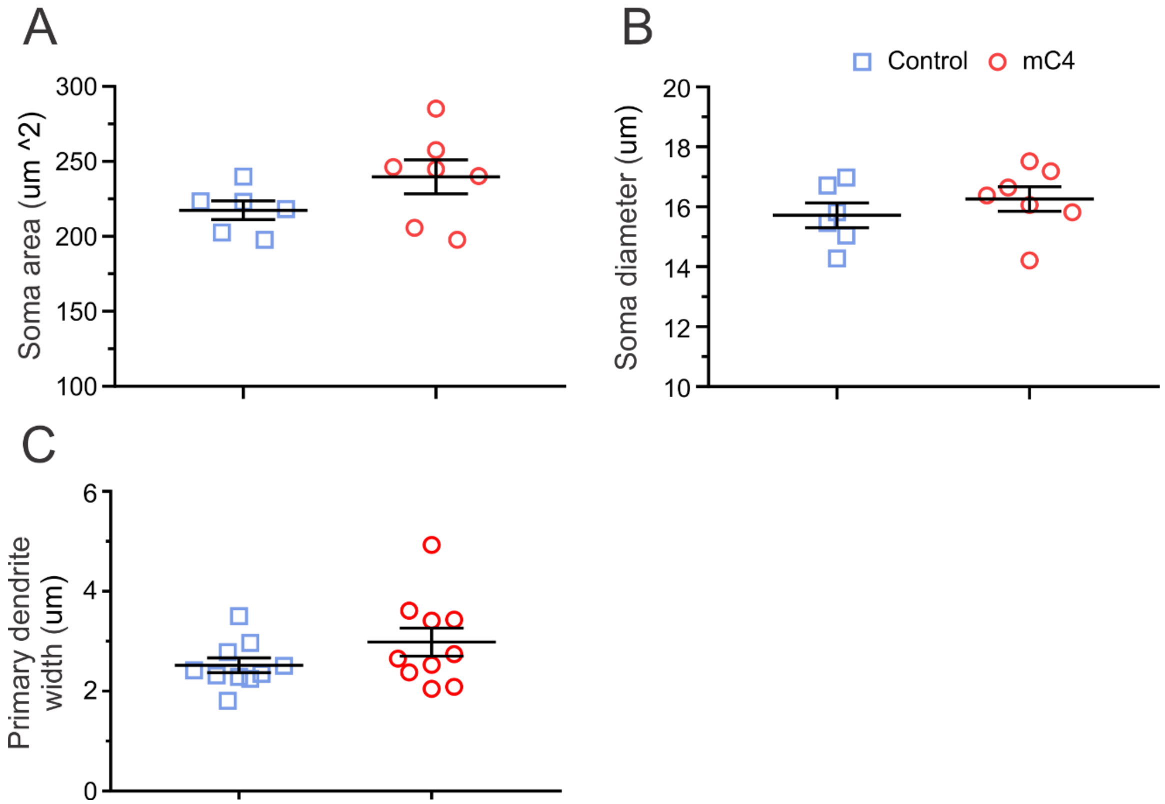


**Supplemental Figure 2: Overexpression of mC4 does not alter soma size or proximal dendrite width.**
**(A)** Soma area is not different between control and mC4 conditions. t-test. $p = 0.13$. **(B)** mC4 overexpression does not alter the diameter of neurons. t-test. $p = 0.37$. **(A-B)** Only GFP-positive L2/3 mPFC neurons included in analysis. Data points represent average measures from ROIs containing many neurons from 3 mice per condition. Control: N = 6 ROIs (including 316 neurons). mC4: N = 7 ROIs (including 216 neurons). **(C)** Primary dendrite width is not different between conditions. N = 10 neurons per condition. Data points represent average primary dendrite width per neuron, including all primary apical and basal dendrites. t-test. $p = 0.16$. Mean ± SEM.

**Supplemental Figure 3**

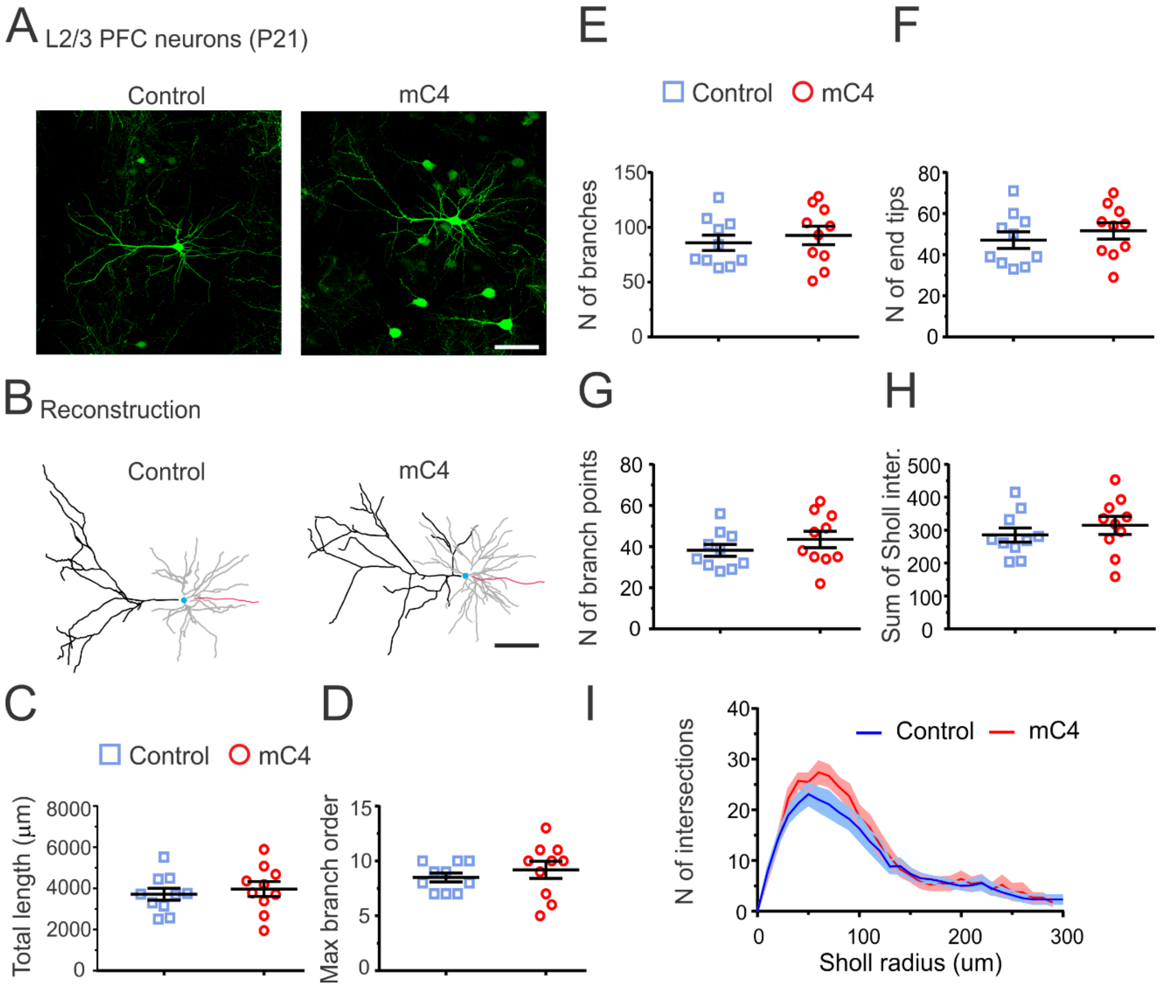


**Supplemental Figure 3: Dendrite morphology is not altered by mC4 overexpression at P21.**
**(A)** Representative confocal images (40X) of control and mC4 GFP-positive L2/3 neurons in the mPFC at P21. Images are max z-projections. Scale bar = 50 µm. **(B)** Reconstructions of control and mC4 neurons from (A). Black lines: apical dendrites. Gray lines: basal dendrites. Red line: axon. Light blue: cell body. Scale bar = 50 µm. **(C)** There is no difference in total dendritic length (µm) between control and mC4 neurons. t-test. *p* = 0.59. **(D)** There is no difference in maximum branch order between control and mC4 neurons. t-test. *p* = 0.44. **(E)** mC4 overexpression does not change the total number of branches in PFC L2/3 neurons. t-test. *p* = 0.54. **(F)** There is no difference in the total number of dendritic end tips between control and mC4 neurons. t-test. *p* = 0.44. **(G)** There is no difference in total number of branch points between conditions. t-test. *p* = 0.29. **(H)** No difference found in the sum of Sholl intersections between control and mC4 neurons. t-test. *p* = 0.41. **(I)** Number of intersections as a function of Sholl radii (µm). Dark blue line: control mean. mC4, Dark red line: mC4 mean. Light blue shade: control SEM. Light red shade: mC4 SEM. **(C-I)** N = 10 neurons per condition. Blue data points: control. Red data points: mC4. Mean ± SEM.

**Supplemental Figure 4**

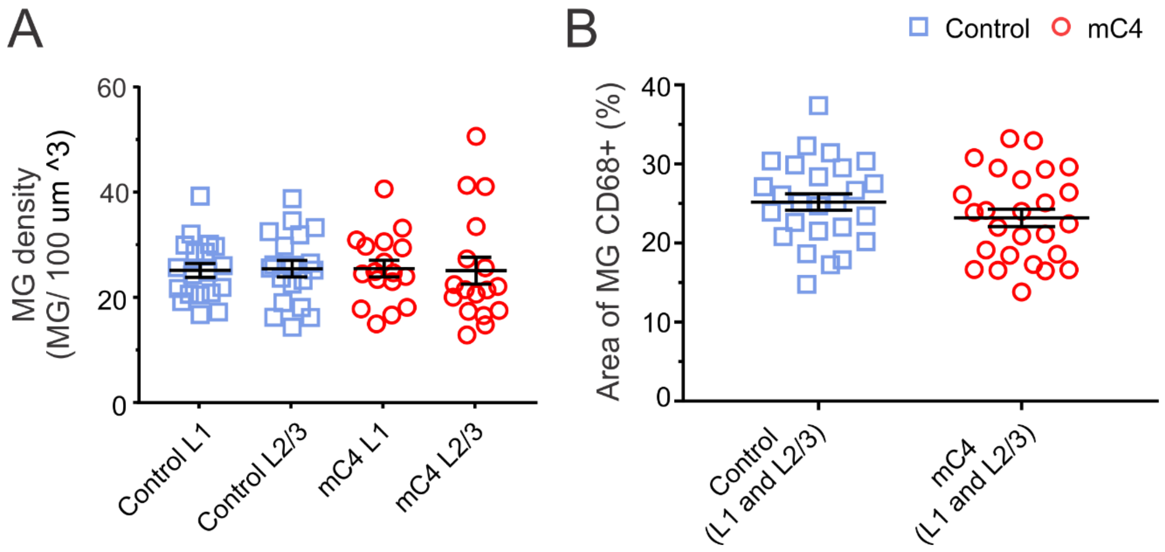


**Supplemental Figure 4: Microglia density and lysosome size are not altered by mC4 overexpression.**
**(A)** Microglia density in superficial layers of the mPFC is not affected by mC4 overexpression. Control: N = 19 ROIs (from 5 mice including 2146 microglia). mC4: N = 17 ROIs (from 5 mice including 1640 microglia). One-way ANOVA with Bonferroni's multiple comparisons. $p$ = 0.998. **(B)** Microglia lysosomal area, as measured by area of MG positive for CD68, is not different between conditions. Area of MG CD68+ (%) = Area of MG CD68+ / Total MG Area. Control: N = 26 ROIs (from 5 mice including 345 microglia). mC4: N =

### Supplemental Figure 5

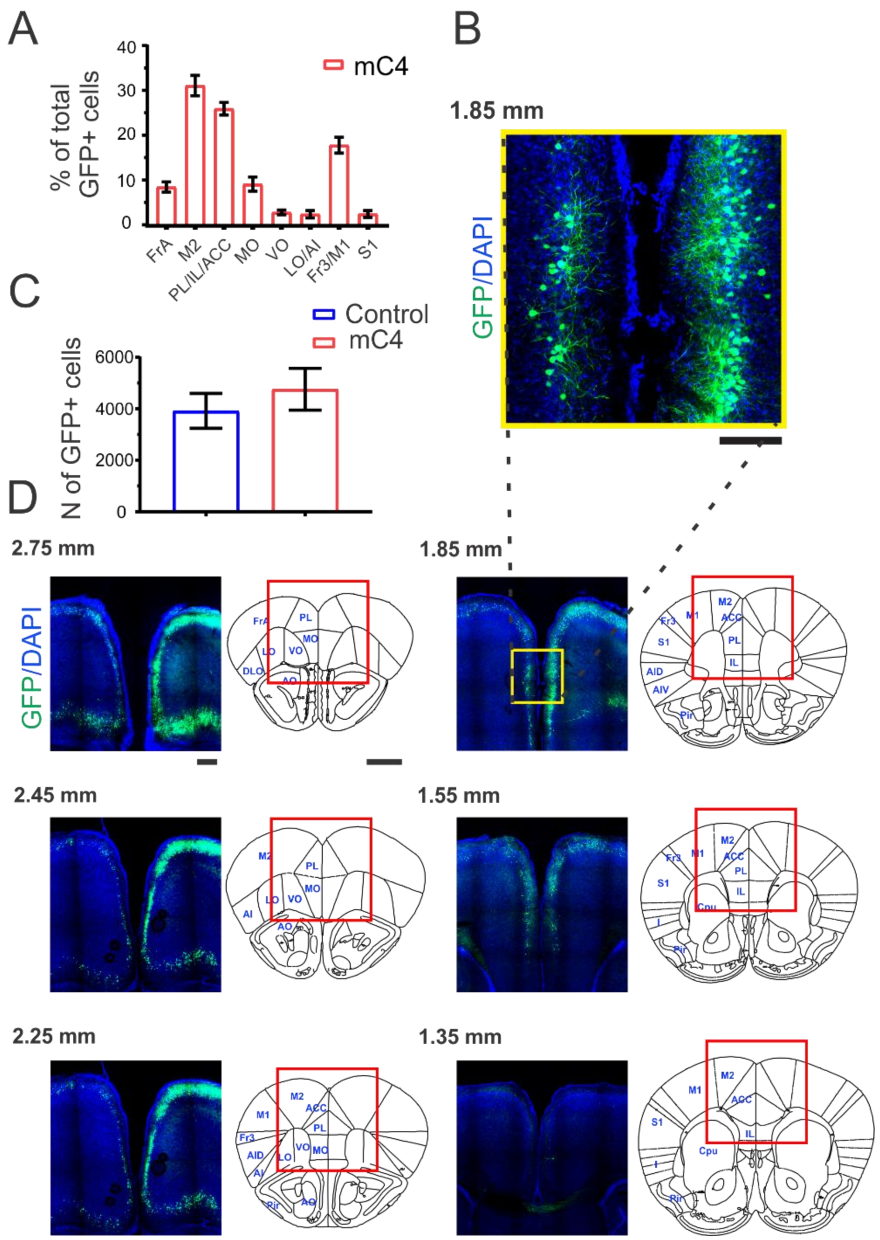


**Supplemental Figure 5: Targeting large populations of L2/3 frontal cortex neurons with *in utero* electroporation.**
**(A)** Percentage of GFP+ cells per area in mC4 mice. N = 21 mC4 mice. **(B)** Representative confocal image (10X) of L2/3 neurons bilaterally co-transfected with mC4 and GFP (green) using IUE. Bregma, +1.85 mm. Scale bar = 150 µm. **(C)** Total number of GFP-positive cells per mouse. N = 36 mice (15 control and 21 mC4). **(D)** Representative sections showing rostro-caudal extent of transfections in the frontal cortex. Images in left panels are zoomed areas from the right panels (red square). Yellow square, zoomed region shown in (B). frontal association cortex (FrA), supplementary motor cortex (M2), prelimbic cortex (PL), infralimbic cortex (IL), anterior cingulate cortex (ACC), medial orbitofrontal cortex (MO), ventral orbitofrontal cortex (VO), lateral orbitofrontal cortex (LO), anterior insular cortex (AI), frontal cortex area 3 (Fr3), primary motor cortex (M1), primary somatosensory cortex (S1), piriform cortex (Pir), anterior olfactory nucleus (AO), caudate-putamen (Cpu). Black numbers: Bregma coordinates. Left panel scale bar = 0.5 mm. Right panel scale bar = 1 mm. Mean ± SEM.

**Supplemental Figure 6**

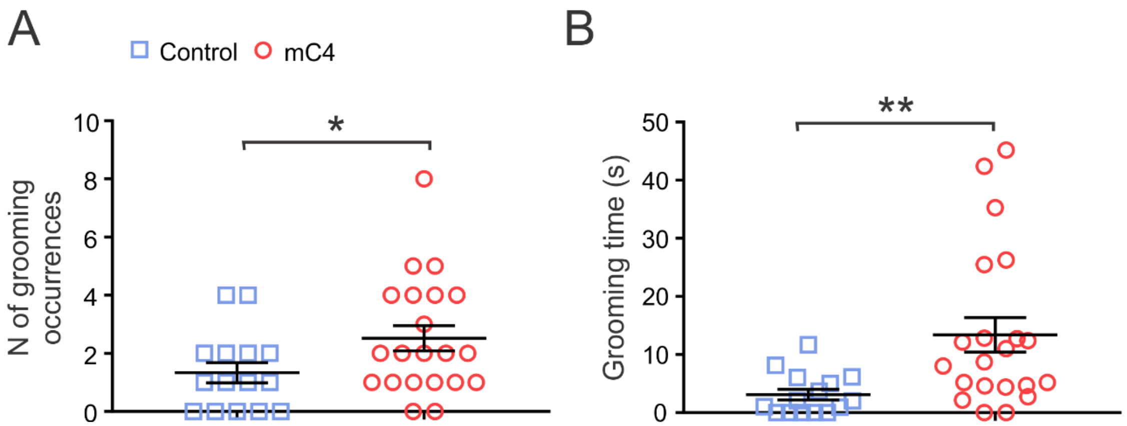


**Supplemental Figure 6: mC4 pups have increased grooming occurrences and grooming time compared to controls.**
**(A)** mC4 mice have greater number of grooming occurrences during the MI 1 task. t-test with Welch's correction. *$p < 0.05$. **(B)** Average time per each grooming occurrence is longer for mC4 mice compared to controls in the MI 1 task. t-test with Welch's correction. **$p < 0.01$. N = 36 mice (15 control and 21 mC4). Mean ± SEM.
